# Cardiac glycosides protect wormseed wallflower (*Erysimum cheiranthoides*) against some, but not all, glucosinolate-adapted herbivores

**DOI:** 10.1101/2023.09.19.558517

**Authors:** Gordon C. Younkin, Martin L. Alani, Anamaría Páez Capador, Hillary D. Fischer, Mahdieh Mirzaei, Amy P. Hastings, Anurag A. Agrawal, Georg Jander

**Author notes:** Present address: Whitehead Institute for Biomedical Research and Department of Biology, Massachusetts Institute of Technology, Cambridge, MA, 02142, USA. Present address: Departamento de Ciencias Biológicas, Universidad de Los Andes, Bogotá, Colombia (Anamaría Páez Capador).

## Abstract

- The chemical arms race between plants and insects is foundational to the generation and maintenance of biological diversity. We asked how the evolution of a novel defensive compound in an already well-defended plant lineage impacts interactions with diverse herbivores. *Erysimum cheiranthoides* (Brassicaceae), which produces both ancestral glucosinolates and novel cardiac glycosides, served as a model.
- We analyzed gene expression to identify cardiac glycoside biosynthetic enzymes in *E. cheiranthoides* and characterized these enzymes via heterologous expression and CRISPR/Cas9 knockout. Using *E. cheiranthoides* cardiac glycoside-deficient lines, we conducted insect experiments in both the laboratory and field.
- *Ec*CYP87A126 initiates cardiac glycoside biosynthesis via sterol side chain cleavage, and *Ec*CYP716A418 has a role in cardiac glycoside hydroxylation. In *Ec*CYP87A126 knockout lines, cardiac glycoside production was eliminated. Laboratory experiments with these lines revealed that cardiac glycosides were highly effective defenses against two species of glucosinolate-tolerant specialist herbivores but did not protect against all crucifer-feeding specialist herbivores in the field. Cardiac glycosides had lesser to no effect on two broad generalist herbivores.
- These results begin elucidation of the *E. cheiranthoides* cardiac glycoside biosynthetic pathway and demonstrate *in vivo* that cardiac glycoside production allows *Erysimum* to escape from some, but not all, specialist herbivores.

## INTRODUCTION

Due to its importance in shaping ecological communities and the species, the chemical arms race between plants and their insect herbivores is a heavily investigated area of molecular ecology (Fraenkel, 1959; Gordon, 1961; Ehrlich & Raven, 1964; Feeny, 1977). Under this paradigm, plants that evolve the ability to produce toxic or deterrent metabolites protect themselves from herbivore feeding and enter a “new adaptive zone” in which they may rapidly diversify in the absence of natural enemies (Ehrlich & Raven, 1964). However, as their enemies evolve the ability to tolerate or neutralize these metabolites, they may in turn enter this protected zone (Gordon, 1961), thereby re-applying ecological pressures that force plants to further adapt their defenses.

The Brassicaceae, a family of more than 4000 plant species, presents many instances of this chemical arms race between plants and specialized herbivores. Glucosinolates evolved as a defense in this lineage approximately 90 million years ago and facilitated multiple rounds of species radiations, resulting in the current high species diversity of the Brassicaceae (Edger *et al*., 2015). Since the original gain of glucosinolate biosynthesis, many insect species have adapted by evolving the ability to tolerate, detoxify, or sequester these compounds (Feeny, 1977; Okamura *et al*., 2022). Several lineages within the Brassicaceae have more recently evolved the production of additional toxic compounds as a second line of defense: globe candytuft (*Iberis umbellata* L.) makes cucurbitacins (Dong *et al*., 2021), garlic mustard (*Alliaria petiolata* Bieb.) makes hydroxynitrile glucosides (Frisch & Møller, 2012), scurvy-grass (*Cochlearia* spp. L.) makes tropane alkaloids (Brock *et al*., 2006), wintercress (*Barbarea vulgaris* W. T. Aiton) makes saponins (Shinoda *et al*., 2002), and wallflowers (*Erysimum* spp. L.) make cardiac glycosides (cardiac glycosides that inhibit Na^+^,K^+^ ATPases in animal cells) (Makarevich *et al*., 1994). It is hypothesized that these key evolutionary innovations allowed these lineages to escape their glucosinolate-tolerant specialized herbivores and again diversify (Thompson, 1989; Züst *et al*., 2020; Dong *et al*., 2021).

A meta-analysis of phytochemical coevolution theory found that specialist insect herbivores are more sensitive than generalists to compounds that are not found in their typical host plants (Cornell & Hawkins, 2003). Consistent with this theory, the glucosinolate-tolerant small and large white cabbage butterflies (*Pieris rapae* L. and *Pieris brassicae* L.) use a wide range of Brassicaceae as host plants but generally avoid *Erysimum*. Experiments involving bioactivity-guided fractionation identified cardiac glycosides as potential agents of this deterrence, and painting cardiac glycosides onto cabbage leaves further established a causal link between the isolated compounds and herbivore behavior (Rothschild *et al*., 1988; Renwick *et al*., 1989; Sachdev-Gupta *et al*., 1993). Similarly, isolated cardiac glycosides were shown to be feeding deterrents for three crucifer-feeding specialist flea beetles (Nielsen, 1978a,b). However, in the absence of mutant lines that lack cardiac glycosides, it has not been possible to conduct *in vivo* tests of the role cardiac glycosides play in *Erysimum*’s escape from herbivory by glucosinolate-tolerant herbivores.

The aims of this study were two-fold: First, we sought to identify cardiac glycoside biosynthesis enzymes in wormseed wallflower (*Erysimum cheiranthoides* L.), in particular the enzyme responsible for catalyzing the first committed step at the branch point between sterol metabolism and cardiac glycoside metabolism, *i.e.*, the cleavage of the sterol side chain to produce pregnenolone (**2**) (Fig. **1**). Second, we revisited classical ecological experiments with modern tools, developing a protocol for stable transformation of *E. cheiranthoides* and generating CRISPR/Cas9-mediated knockouts of cardiac glycoside biosynthetic genes. This effectively reversed the “escape from herbivory” of *E. cheiranthoides* and allowed us to directly test the hypothesis that cardiac glycosides in *Erysimum* provide a targeted defense against crucifer-specialist herbivores, both in the laboratory and in the field.

**Fig. 1.**
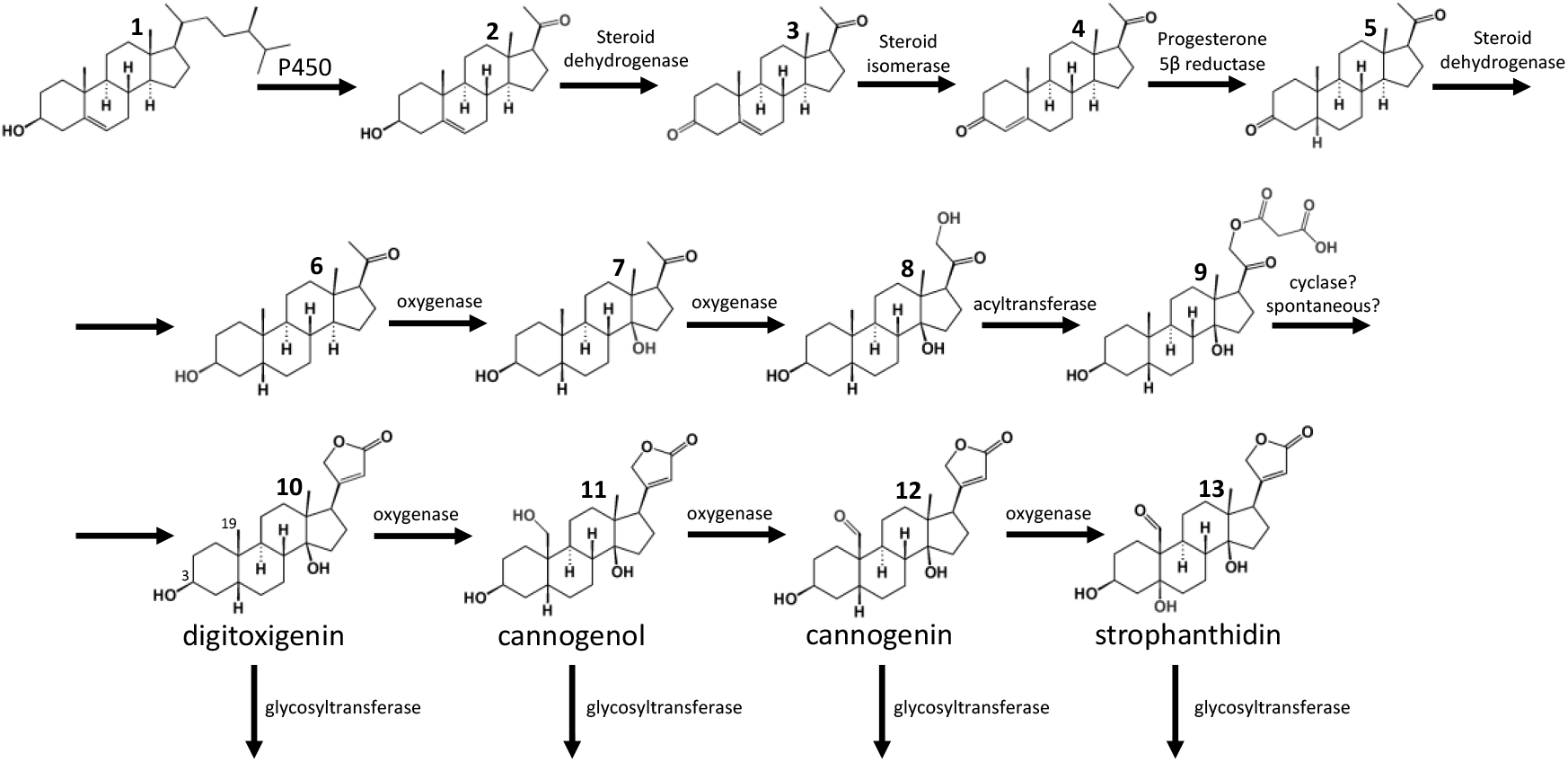
Proposed *Erysimum* cardiac glycoside biosynthetic pathway. The first step (sterol side chain cleavage), and the conversion of digitoxigenin **10** to cannogenol **11** are discussed in this paper. Relevant carbon numbering is shown for digitoxigenin. One or two sugars may be attached at the 3-hydroxyl group.

## MATERIALS AND METHODS

### Plants, insects, and growth conditions

All experiments were conducted with the genome-sequenced *E. cheiranthoides* var. Elbtalaue, which has been inbred for at least eight generations (Züst *et al*., 2020), Arabidopsis Biological Resource Center (https://abrc.osu.edu) accession number CS29250. Plants were grown in Cornell Mix (by weight 56% peat moss, 35% vermiculite, 4% lime, 4% Osmocote slow-release fertilizer [Scotts, Marysville, OH], and 1% Unimix [Scotts]) in Conviron (Winnipeg, CA) growth chambers with a 16:8 photoperiod, 180 μM m^−2^ s^−1^ photosynthetic photon flux density, 60% humidity, and constant 23 °C temperature. Once flowering, plants were moved to a climate-controlled greenhouse set to 26 °C during the day and 24 °C at night. Natural light was supplemented with artificial light set to a 16:8 hour photoperiod.

Cabbage looper (*Trichoplusia ni* Hübner) eggs were obtained from Benzon Research (Carlisle, PA) and hatched on artificial diet (Southland Products, Lake Village, AR) in an incubator at 28 °C. Wild-caught *P. rapae* butterflies (Ithaca, NY, USA, June 2023) were used to start a lab colony. Adults were fed a 10% sucrose solution and were presented with *Brassica oleracea* var. capitata for oviposition and caterpillar feeding. Green peach aphids (*Myzus persicae* Sulzer) were from a lab colony of a previously described, genome-sequenced “USDA” strain (Ramsey *et al*., 2007, 2014; Feng *et al*., 2023). Cabbage aphids (*Brevicoryne brassicae* L.) were from a colony originally collected in 2015 by Brian Nault (Cornell University) in Geneva, NY, USA. Both aphid species were maintained on *B. oleracea* var. capitata in a growth room with a 16:8 photoperiod and constant 23 °C temperature.

### RNA-sequencing analysis

Raw RNA sequencing reads from 48 *Erysimum* species (Züst *et al*., 2020) were downloaded from the NCBI Short Read Archive (PRJNA563696)(Strickler *et al*., 2019). Tissue-specific RNA sequencing data were collected from six-week-old wildtype *E. cheiranthoides* plants, including roots and young leaves that had just emerged, measuring 1 cm in length (PRJNA1015726). The SV Total RNA Isolation System (Promega Corporation, Madison, WI, USA) was employed to isolate total RNA, and the quality and quality of RNA were evaluated using the RNA Integrity Number (RIN) from a 2100 Bioanalyzer (Agilent Technologies, Santa Clara, CA). For sequencing, 5 μg of purified total RNA, pooled from three replicates, was used for the preparation of strand-specific RNAseq libraries with 14 cycles of final amplification (Zhong *et al*., 2011). The libraries were multiplexed and sequenced with a paired-end read length of 150 bp using two lanes on an Illumina HiSeq2500 instrument (Illumina, San Diego, CA) at the Cornell University Biotechnology Resource Center (Ithaca, NY).

Raw RNA sequencing reads for species and tissue datasets were pseudoaligned to the *E. cheiranthoides* genome v2.0 (PRJNA563696)(Strickler *et al*., 2019) using kallisto with default parameters, yielding transcript counts (Bray *et al*., 2016). Output files were normalized and transformed using the transform_counts.R script from the mr2mods pipeline (Wisecaver et al., 2017; https://github.itap.purdue.edu/jwisecav/mr2mods). Fold-change expression between leaf and root tissue was calculated using edgeR (Robinson *et al*., 2010; McCarthy *et al*., 2012).

### Cloning of candidate genes

*Erysimum cheiranthoides* RNA was extracted from 2-week-old seedlings and young leaves of 5- week-old plants using the SV Total RNA Isolation System, and cDNA was generated using SMARTScribe Reverse Transcriptase (Takara Bio USA, Ann Arbor, MI). Primers were ordered to include Gateway *attB* recombination sites (Table **S1**), and the coding sequence was amplified from cDNA using Phusion High-Fidelity DNA Polymerase (New England Biolabs, Ipswich, MA). The gel-purified amplicon was inserted into the pDONR207 vector using Gateway BP Clonase II enzyme mix (ThermoFisher Scientific, Waltham, MA) and then into pEAQ-HT-DEST1 (Sainsbury *et al*., 2009) using Gateway LR Clonase II enzyme mix (ThermoFisher Scientific). The sequences of the inserted genes were verified with Sanger sequencing. All cloning was done using 10-beta Competent *E. coli* (NEB, Ipswich, MA), with transformations using heat shock at 42 °C. Plasmids were purified using the Wizard Plus SV Miniprep DNA Purification System (Promega Corporation) and transformed into *Agrobacterium tumefaciens* strain GV3101 using a freeze-thaw method (Weigel & Glazebrook, 2006).

### *Transient expression of candidate genes in* Nicotiana benthamiana

Genes were transiently expressed in leaves of 4-week-old *Nicotiana benthamiana* plants (Bach *et al*., 2014). A single colony of *A. tumefaciens* strain GV3101 carrying pEAQ-HT-DEST1 containing the candidate gene was inoculated into a 10 mL culture of lysogeny broth, pH 7.5 (LB), with 50 μg/mL rifampicin, 20 μg/mL gentamicin, and 50 μg/mL kanamycin and shaken for 24 hours at 28 °C and 230 rpm. The bacteria were pelleted for 10 minutes at 3200 rcf in an Eppendorf Centrifuge 5810 (Hamburg, Germany) and resuspended to OD_600_=0.5 in a solution containing 10 mM 2-(N-morpholino)ethanesulfonic acid (MES), 10 mM MgCl_2_, and 400 μM acetosyringone before resting in the dark for 2 hours prior to infiltration into the abaxial surface of leaves using a blunt syringe. Each construct was infiltrated into leaves of at least three separate plants, with pEAQ-HT-DEST1 carrying GFP serving as a negative control. In the case of *Ec*CYP716A418, for which the substrates were not predicted to be present in *N. benthamiana* leaves, a 200 μM solution of digitoxigenin (Sigma-Aldrich, St. Louis, MO) or digitoxin (Sigma-Aldrich) was infiltrated three days after infiltration with *A. tumefaciens*. Tissue was collected five days after *A. tumefaciens* infiltration for UPLC-MS analysis. A 200 μM solution of pregnenolone (Sigma-Aldrich) was also infiltrated into separate *N. benthamiana* leaves two days prior to tissue harvest to check for any modifications that may occur *in planta*.

### gRNA design and CRISPR/Cas9 constructs

One or two CRISPR guide RNAs (gRNAs) were designed to target the first exon of each candidate gene using the IDT CRISPR-Cas9 guide RNA design tool. Single-stranded DNA oligos were ordered for each gRNA, one containing the forward gRNA sequence and a 5’ ATTG, and one containing the reverse complement and a 5’ AAAC (Table **S1**). Complementary oligos were annealed and inserted into either pARV483 in the case of a single gRNA or into pARV370 in the case of multiple gRNAs targeting the same gene, using Type IIS restriction enzyme Aar1 (New England Biolabs). gRNA cassettes including the AtU6-26 promoter, gRNA scaffold, and AtU6-26 terminator were PCR amplified from pARV370 using primers containing PaqCI (New England Biolabs) restriction sites (Table **S1**) and inserted in tandem into pARV380 such that all gRNAs targeting the same gene were contained on a single plasmid. Plasmid maps for pARV483, pARV370, and pARV380 are provided in Fig. **S1**.

### *Floral dip stable transformation of* Erysimum cheiranthoides

A floral dip stable transformation protocol for *E. cheiranthoides* was developed based on methods previously published for *Arabidopsis thaliana* (Clough & Bent, 1998) and *Brassica napus* (Wang *et al*., 2003). *Agrobacterium tumefaciens* strain GV3101 containing a binary plasmid was grown overnight as described above for *N. benthamiana* assays, but the initial 10 mL culture was inoculated into 200 mL fresh LB with the same antibiotics and cultured under the same growth conditions for an additional 24 hours.

To prepare the infiltration solution, *Agrobacterium* cultures were spun down at 3200 rcf in a Eppendorf Centrifuge 5810 for 10 minutes at room temperature and resuspended in a solution containing full strength Murashige and Skoog (MS) salts (Research Products International, Mt. Prospect, IL), 50 g/L sucrose (Sigma-Aldrich), 0.1 mg/L 6-benzylaminopurine (Sigma-Aldrich), 400 μM acetosyringone (Sigma-Aldrich), and 0.01% Silwet L-77 (PlantMedia.com, Chiang Mai, Thailand) and were allowed to rest for one hour. The inflorescence of *E. cheiranthoides* plants just beginning to flower was submerged in the bacterial suspension, agitated, and placed under vacuum for 5 minutes. The vacuum was quickly released, and inflorescence was covered with plastic wrap and secured with a twist tie. Plants were kept in dark for 18-24 hours before removing the plastic wrap and transferring to greenhouse conditions. Seeds were harvested 6 weeks after dipping to be screened for transformants by looking for *DsRed* fluorescence using an SZX12 stereomicroscope equipped with a UV lamp (Olympus, Center Valley, PA).

### T7 Endonuclease 1 assay for detecting Cas9-induced mutations

T1 plants were screened for mutations using a T7 endonuclease 1 (T7E1) assay. DNA was extracted from 3-week-old T1 plants by heating a 1 mm leaf disk in 25 μL Extract-N-Amp extraction solution (E7526) at 95 °C for 10 minutes and then adding 25 μL PCR Diluent (E8155, MilliporeSigma, St. Louis, MO). Primers were selected to amplify an approximately 1000 bp region flanking the Cas9 target site (Table **S1**). PCR was carried out using Phire Green Hot Start II Mastermix (ThermoFisher Scientific, Waltham, MA) under manufacturer-recommended conditions. For the T7E1 assay, the Alt-R Genome Editing Detection Kit (Integrated DNA Technologies, Coralville, IA) was used according to manufacturer specifications. In any samples with the presence of non-wildtype amplicons, PCR products were purified (Wizard SV Gel and PCR Clean-Up System, Promega Corporation, Madison, WI) and sent for Sanger sequencing at the Cornell Biotechnology Resource Center (Cornell University, Ithaca, NY). T2 seeds collected from plants with confirmed target site mutations were screened for the absence of fluorescence by microscopy and for presence of a homozygous mutation at the target site using Sanger sequencing. T3 seeds collected from these T-DNA-free, homozygous mutant plants were used for further analyses.

### *Metabolite feeding to cyp87a126* E. cheiranthoides *mutants*

Predicted cardiac glycoside intermediates were fed to *cyp87a126-2* mutant plants to check for rescue of cardiac glycoside biosynthesis. Two hundred μM pregnenolone, isoprogesterone (TLC Pharmaceutical Standards, Newmarket, ON), progesterone (Sigma-Aldrich), or 5β-pregnane-3,20-dione (aablocks, San Diego, CA) were suspended in 10 mM MES, 10 mM MgCl_2_ and injected into the abaxial surface of young leaves of 4-week-old plants. Tissue was collected after 2 days for UPLC-MS analysis.

### Metabolite extraction from plant tissue

The same metabolite extraction protocol was used for tissue of both *E. cheiranthoides* and *N. benthamiana*. Two 14 mm leaf disks were collected into a 1.7 mL microcentrifuge tube (Laboratory Products Sales Inc., Rochester, NY, USA), either from an infiltrated region of leaf in the case of infiltration experiments, or from the youngest fully expanded leaves of four to five-week-old plants in the case of all other experiments. Tissue was flash frozen in liquid nitrogen and ground with three 3-mm ball bearings (Abbott Ball Company, Hartford, CT) on a 1600 MiniG^TM^ tissue homogenizer (SPEX SamplePrep, Metuchen, NJ). One hundred μL 70% methanol with 15 μM internal standard was added to each sample (ouabain for positive ionization mode, sinigrin for negative ionization mode), which was then vortexed to suspend the plant tissue. Samples were left to extract for half an hour at room temperature before being centrifuged for ten minutes at 17,000 rcf in a Z207-M microcentrifuge (Hermle, Sayreville, NJ). The supernatant was transferred to a clean 1.7 mL tube and centrifuged again for ten minutes at 17,000 rcf before being transferred to vials for UPLC-MS analysis.

### Ultrahigh pressure liquid chromatography coupled to mass spectrometry (UPLC-MS)

Plant and yeast extracts were analyzed on an UltiMate 3000 UHPLC system coupled to a Q-Exactive hybrid quadrupole-orbitrap mass spectrometer (ThermoFisher Scientific). The instrument was fitted with a Supelco Titan^TM^ C18 UHPLC Column (80Å, 100 x 2.1 mm, particle size 1.9 μm; Sigma Aldrich). Two μL injections were separated using a short (for quantification) or long (for figures) solvent gradient. Mobile phase A was water + 0.1% (v/v) formic acid and mobile phase B was acetonitrile + 0.1% (v/v) formic acid. All solvents were Optima LC/MS grade (Thermo Fisher Scientific). Short gradient: 0-0.5 minutes, hold at 2% B; 0.5-10 minutes, linear gradient from 2%-97% B; 10-11.5 minutes, hold at 97% B, 11.5-13 minutes, hold at 2% B. Long gradient: 0-5 minutes, hold at 2% B; 5-22 minutes, linear gradient from 2%-97% B; 22-23.5 minutes, hold at 97% B, 23.5-25 minutes, hold at 2% B. The solvent flow rate was 0.5 mL/minute, the column oven was set to 40 °C, and the autosampler temperature was 15 °C for all methods. Mass spectrometry data were acquired in positive ionization mode for detection of cardiac glycosides and negative ionization mode for detection of glucosinolates in full scan mode from m/z 150-900. The following settings were used: capillary voltage 3.5 kV (positive mode) or 3.0 kV (negative mode), capillary temperature 380 °C, sheath gas 60, auxillary gas 20, and S-Lens RF level 50.

### LCMS data processing

For targeted LC-MS analysis, peak areas were quantified using a custom processing method in Xcalibur^TM^ Software (Thermo Fisher Scientific) using the following parameters: peak detection ICIS, smoothing points 1, baseline window 40, area noise factor 5, peak noise factor 15, tailing factor 2. Mass features used for quantification are provided in Table **S2** for cardiac glycosides and Table **S3** for glucosinolates.

### Na^+^,K^+^-ATPase Inhibition Assay

The inhibitory effect of plant extracts on porcine (*Sus scrofa* L) Na^+^,K^+^-ATPase was measured following the protocol described in Petschenka et al. (Petschenka *et al*., 2023). Wildtype *E. cheiranthoides* was compared with *E. cheiranthoides* mutant lines *cyp87a126-1* and *cyp87a126- 2*, and *A. thaliana* ecotype Columbia (Col-0) served as a glucosinolate-containing, cardiac glycoside-free control. Two biological replicates of wildtype *E. cheiranthoides* (each with two technical replicates) and four biological replicates of *cyp87a126-1*, *cyp87a126-2*, and *A. thaliana* (each with one technical replicate) were assayed. Using the enzymatic activity across sample dilutions, sigmoid dose-response curves were calculated using a logistic function in the nlme package (Pinheiro & Bates, 2000, 2023) in R statistical software (R Core Team, 2020). For each sample, the relative dilution at the inflection point was calculated to estimate the half maximal inhibitory concentration (IC50).

### Insect Bioassays

For caterpillar growth and survival assays, individual 2-day-old *T. ni* or *P. rapae* larvae were restricted to a single leaf of 4-week-old of *E. cheiranthoides* wildtype, *cyp87a126-1*, and *cyp87a126-2* mutant lines (Fig. **S2**) using 6.5 × 8 cm organza bags (amazon.com, item B073J4RS9C). For *T. ni*, 12 plants were used for each line, and caterpillars were placed on five leaves of each plant for a total of 60 caterpillars per line. Some caterpillars were removed due to underwatering of plants or caterpillar death/escape (WT: 37 caterpillars removed, *cyp87a126-1*: 25, *cyp87a126-2*: 32). For *P. rapae*, a variable number of plants per line was used (WT: 14 plants, *cyp87a126-1*: 17 plants, *cyp87a126-2*: 13 plants), with one caterpillar on a single leaf of the same age per plant. Caterpillars were. After eight days, leaf damage was assessed, and surviving larvae were moved to a fresh leaf to continue feeding.

Aphid and caterpillar choice assays were conducted in 100×15 mm Petri dishes (ThermoFisher Scientific) sealed with Parafilm. For *T. ni*, 14 mm leaf disks from young leaves were placed in pairs of one wildtype and one mutant leaf disk on wet paper towels along with a single neonate caterpillar, with 20 replicates per mutant line, for a total of 40 replicates (Fig. **S2**). After 48 hours, photos were taken of each leaf disk, and leaf area eaten was quantified using the Leaf Byte app (Getman-Pickering *et al*., 2020). Two replicates for each line were removed due to desiccation of leaves. For *M. persicae* and *B. brassicae*, detached leaves were used instead of leaf disks, and 10 adult aphids were placed in each Petri dish (Fig **S2**). Twelve replicates were assayed for each mutant line, and after 24 hours, the number of aphids on each leaf was recorded. Any replicates for which either or both disks were shriveled or limp at the end of the experiment were removed (*M. persicae*: two replicates removed for each line, *B. brassicae*: three removed for *cyp87a126-1*, one removed for *cyp87a126-2*).

*Myzus persicae* and *B. brassicae* colony growth was measured by transferring five adult aphids from cabbage to bagged 3-week-old *E. cheiranthoides* plants, with 12 replicates for wildtype and each mutant line. After 9 days, the combined number of adults and nymphs was recorded for each plant.

*Pieris rapae* oviposition assays were conducted using lab-reared adult butterflies. One wildtype plant and one mutant plant (either *cyp87a126-1* or *cyp87a126-2*) were placed in a 38 x 38 x 60 cm mesh cage with a mating pair of *P. rapae* butterflies and a 10% sucrose solution (Fig. **S2**). Butterflies were monitored daily and the total number of eggs on each plant was recorded once the female butterfly died or after five days. There were eleven planned replicates for each mutant line, but many butterflies died before laying any eggs, leaving only 6 replicates for *cyp87a126-1* and 4 replicate for *cyp87a126-2*. Because *P. rapae* lay eggs one at a time and not in clusters, replicates were pooled and the distribution of eggs between wildtype and mutant lines was used for statistical analysis.

### Field experiment

Three-week-old *E. cheiranthoides* plants were transplanted to a freshly plowed field in Ithaca, NY, USA on July 31, 2023. Plants were arranged in blocks of six, with each block containing two wildtype plants, one plant each from *cyp87a126-1* and *cyp87a126-2* mutant lines, and two plants from glucosinolate knockdown lines that are otherwise not discussed in this study. Each block was arranged in a 2×3 pattern, with genotypes randomized within each block, and plants spaced 40 cm apart. There were 3 rows of ten blocks spaced 1.1 m apart on all sides, for a total of 30 blocks and 180 plants. There was a 2 m buffer on all sides between the experimental plot and unplowed areas. Visitors to each plant were recorded during four separate censuses over the course of a month and visually identified to the lowest possible taxon. Plants that died before the end of the experiment or that were cut down at ground level by a suspected mammalian herbivore were removed, leaving 52 wildtype, 27 *cyp87a126-1*, and 24 *cyp87a126-2* plants. Only taxa found on at least 25 plants (>24% of surviving plants) across the course of the experiment were retained for statistical analysis.

### Statistical analysis

All statistical analyses were carried out in R statistical software (R Core Team, 2020). The following functions and packages were used: edgeR (Robinson *et al*., 2010; McCarthy *et al*., 2012) for differential gene expression analysis, aov and TukeyHSD for one-way ANOVA and post-hoc Tukey’s HSD for most metabolite abundance and insect assays. A Kruskal-Wallis test (kruskal.test) with pairwise Wilcoxon post-hoc tests (pairwise.wilcox.test) were used for the non-normally distributed results of the *B. brassicae* colony growth assay. For analysis of results from the field experiment, a generalized linear model with a binomial distribution was fitted to presence/absence of each visitor on a given plant across all four censuses using the glm function in base R. In the case of a significant genotype effect, pairwise contrasts and Tukey-adjusted p-values were estimated using emmeans (Lenth, 2023). *Phyllotreta striolata* (F.) was found on every plant at some point, so a model based on presence/absence data was not informative. Instead, a one-way ANOVA (aov function in R) was performed on counts of *P. striolata* beetles per plant. Plots were made using the packages genemodel (Monroe, 2017), MSnbase (Gatto & Lilley, 2012; Gatto *et al*., 2020), multcompView (Graves *et al*., 2023). R scripts for all statistical analyses are available on GitHub (https://github.com/gordonyounkin/EcCYP87A126_scripts), and raw data underlying all figures are available in the Supplementary Information.

### Phylogenetic Inference

Homologous sequences from selected species were identified using BLAST against public databases and were aligned using ClustalW (Sievers *et al*., 2011; Madeira *et al*., 2022). Gene phylogenies were inferred using IQ-TREE web server (Trifinopoulos *et al*., 2016; Hoang *et al*., 2018; Minh *et al*., 2020) with default parameters, except the number of bootstrap alignments was increased to 10,000.

## RESULTS

### Identification of candidate genes for cardiac glycoside biosynthesis

To identify cytochrome P450 monooxygenases involved in cardiac glycoside biosynthesis, we examined patterns of gene expression across *E. cheiranthoides* tissues and between 48 different species in the genus *Erysimum.* Two criteria were used: (1) High expression in leaves of *E. cheiranthoides* relative to roots, as grafting experiments showed that cardiac glycosides are synthesized in leaves and transported to the roots (Alani *et al*., 2021). (2) Lower expression in *Erysimum collinum* relative to all other species of *Erysimum*, because *E. collinum* produces nearly undetectable levels of cardiac glycosides (Züst *et al*., 2020). Of the 116 cytochrome P450s identified across the two expression datasets, only 3 matched both criteria, with at least 9-fold greater expression in young leaves relative to roots, and expression in *E. collinum* leaves more than 3 standard deviations below the mean of expression levels in other *Erysimum* species (Fig. **2a**). These 3 cytochrome P450s, *Ec*CYP71B132, *Ec*CYP716A418, and *Ec*CYP87A126, were selected as candidates for involvement in cardiac glycoside biosynthesis. Full length coding sequences for candidate genes are provided in Table **S4**.

**Fig. 2.**
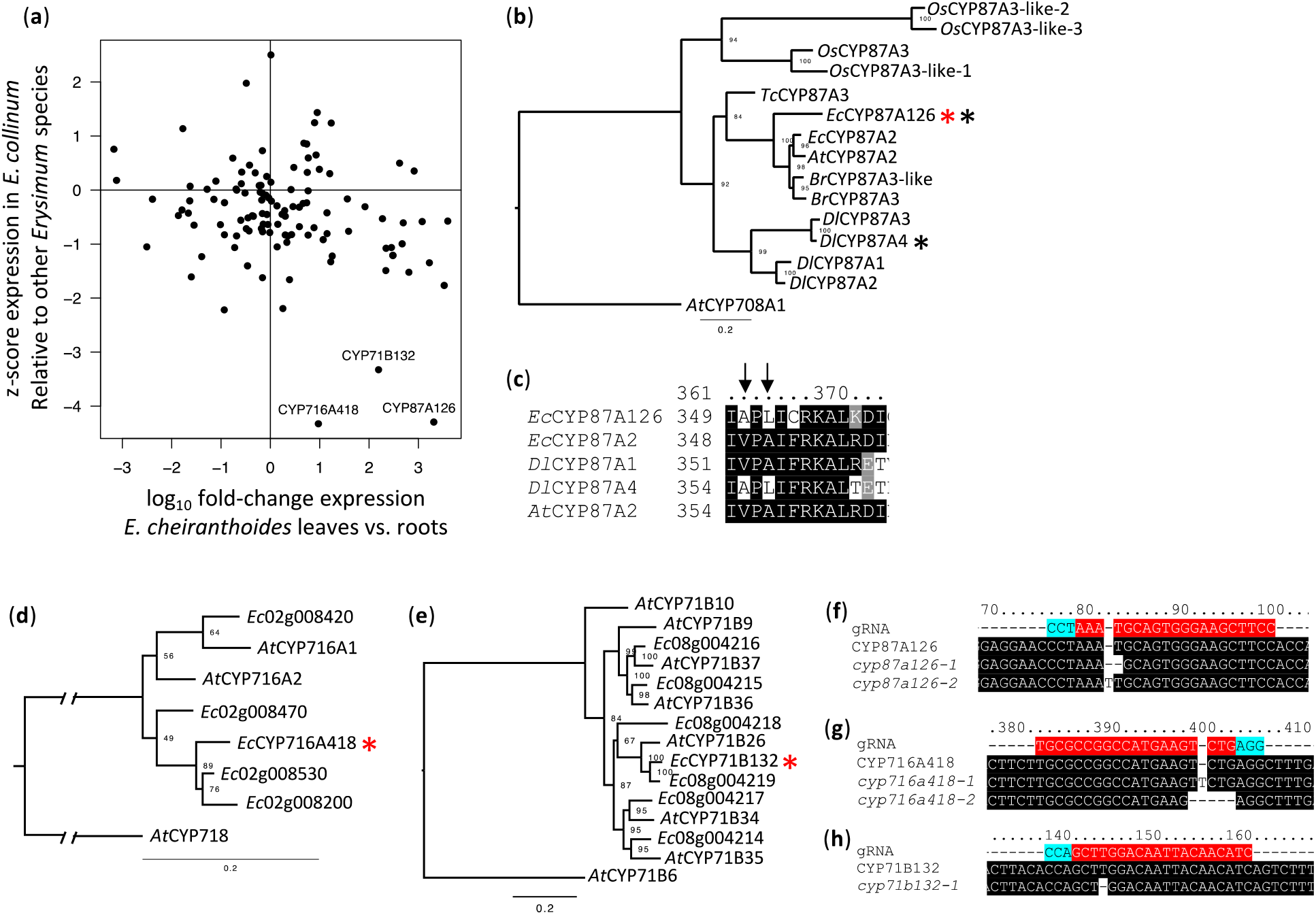
Identification and knockout of candidate cytochrome P450s. (a) Expression pattern of cytochrome P450 monooxygenases in *Erysimum.* Fold change in expression between young leaves and roots is plotted against z-score of expression in *E. collinum* relative to 48 *Erysimum* species. Genes with high expression in young leaves, where cardiac glycosides are synthesized, and low expression in *E. collinum*, which does not produce cardiac glycosides, are good candidates for involvement in cardiac glycoside biosynthesis. (b,d,e) Gene trees of candidate cytochrome P450s. Species abbreviations: *At* (*Arabidopsis thaliana*), *Br* (*Brassica rapa*), *Dl* (*Digitalis lanata*), *Ec* (*Erysimum cheiranthoides*), *Os* (*Oryza sativa*), and *Tc* (*Theobroma cacao*). Candidate genes are marked with a red star. Black stars indicate genes that have been previously shown to have sterol side chain cleaving activity. (c) Alignment of selected CYP87A proteins. Convergent amino acid substitutions that are critical for sterol side chain cleaving activity are marked with an arrow. (f,g,h) Location and sequence of Cas9 protospacers (red, with 3’ NGG PAM in turquoise) used for generation of mutant lines with wildtype and mutant sequences. Number of basepairs from the start of the coding sequence are indicated.

### Phylogenetic analysis of candidate genes

All three candidate genes are in clades containing duplication events relative to *A. thaliana* (Fig. **2b,d****,e**). *Ec*CYP87A126 is of particular interest because of recent reports identifying CYP87A members as capable of sterol side chain cleavage in several species including woolly foxglove (*Digitalis lanata* Ehrh.) (Carroll *et al*., 2023), as well as common foxglove (*D. purpurea* L.), Sodom apple (*Calotropis procera* W. T. Aiton), and *E. cheiranthoides* (Kunert *et al*., 2023). To better understand the convergence of this activity in diverse lineages, we aligned the amino acid sequences of *Ec*CYP87A126 and *Dl*CYP87A4. We found that the two amino acid substitutions identified by Carroll et al. (Carroll *et al*., 2023) as necessary for sterol side chain cleaving activity, V355A and A357L in *Dl*CYP87A4, were also present in *Ec*CYP87A126 (Fig. **2c**)

### *Cardiac glycoside content is altered in Cas9-generated* cyp87a126 *and* cyp716a418 *mutant lines*

We generated independent knockout lines for each of the three cytochrome P450s using Cas9-mediated gene editing (Figs. **2f,g****,h**, **S3**). The knockout lines did not display an obvious visual phenotype (Fig. **3b**), and the *Ec*CYP71B132 knockout line displayed no changes in cardiac glycoside content (Fig. **3a**). Knockout lines for the other two candidate genes had strong alterations in cardiac glycoside accumulation. *Ec*CYP716A418 knockout lines hyperaccumulate glycosides of digitoxigenin (**10**) (Fig. **3a,d**, Table **S5**, one-way ANOVA: F_2,10_=74.01 *P<*0.001; Tukey’s HSD: WT-*cyp716a418-1 P<*0.001, WT-*cyp716a418-2 P<*0.001), apparently lacking the ability to hydroxylate digitoxigenin at carbon 19 to form cannogenol (**11**), cannogenin (**12**), and strophanthidin (**13**) (Fig. **1**). *Ec*CYP87A126 knockout lines accumulated barely detectable levels of cardiac glycosides (Fig. **3a**), suggesting that it is an essential enzyme in cardiac glycoside biosynthesis. Despite a 1000-fold decrease in total cardiac glycoside-related peak area (Fig. **3c**, Table **S6**, one-way ANOVA: F_2,12_=271 *P* <0.001; Tukey’s HSD: WT-*cyp87a126-1 P* <0.001, WT-*cyp87a126-2 P* <0.001), *cyp87a126* lines display no difference in aliphatic (one-way ANOVA: F_2,10_=2.32 *P* =0.15) or indole (one-way ANOVA: F_2,10_=1.71 *P* =0.23) glucosinolate abundance (Fig. **3c**, Table **S7**).

**Fig. 3.**
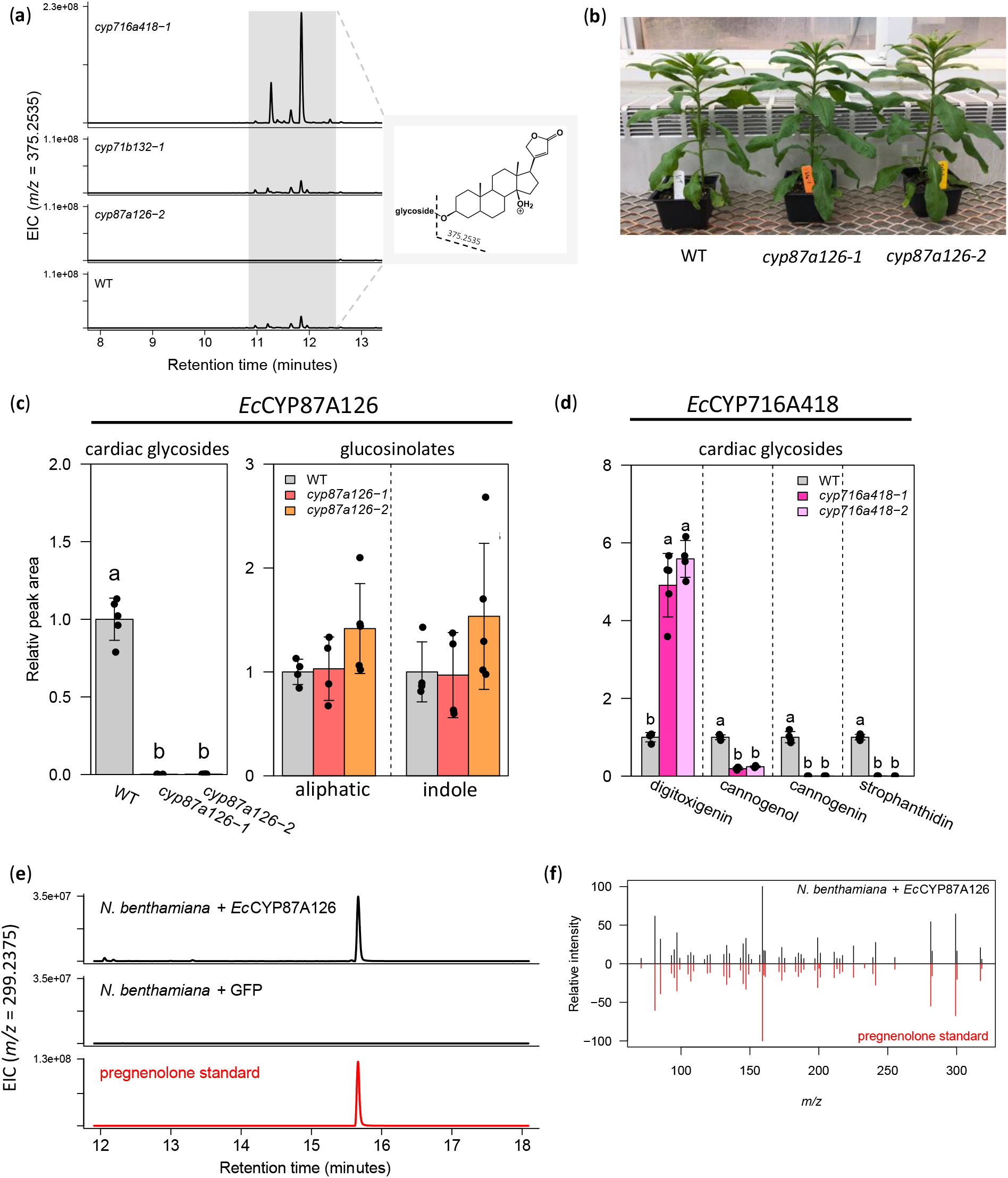
*Ec*CYP87A126 and *Ec*CYP716A418 mutant lines have altered cardiac glycoside content. (a) Extracted ion chromatograms from a representative mutant plant for each candidate gene. *m/z* = 375.2535 is a fragment common to all digitoxigenin-containing cardiac glycosides in positive electrospray ionization. (b) Despite lacking cardiac glycosides, *cyp87a126* mutant plants show no obvious growth phenotype. (c) Total cardiac glycoside and glucosinolate-related peak area in *cyp87a126* mutants compared to wildtype. (d) Cardiac glycoside abundance by genin in *cyp716a418* mutant lines compared to wildtype. For all plots: error bars indicate ± SD (n=4-5); letters indicate statistical differences (*P*<0.001) according to a one-way ANOVA followed by Tukey’s HSD. n=4-5 plants per line. (e) Extracted ion chromatograms (*m/z* = 299.2375, pregnenolone [M-H2O+H]+) of *N. benthamiana* leaves expressing EcCYP87A126 or a GFP control. A pregnenolone standard (red) was infiltrated into a separate *N. benthamiana* leaf to account for any potential modifications made by endogenous enzymes. (f) MSMS spectra of pregnenolone standard (red) compared to product of *Ec*CYP87A126 (black).

### *Transient expression of* E. cheiranthoides *cytochrome P450s in* N. benthamiana *leaves*

To investigate the enzymatic activity of *Ec*CYP716A418 and *Ec*CYP87A126, full length coding sequences were cloned and transiently expressed in *N. benthamiana* leaves, with substrate co-infiltration where necessary. Based on the strong phenotype of the knockout lines, we expected *Ec*CYP716A418 to hydroxylate digitoxigenin at carbon 19. However, no activity was detected upon co-infiltration with digitoxigenin or digitoxin, a glycosylated form of digitoxigenin. *Nicotiana benthamiana* leaves expressing *Ec*CYP87A126 accumulated pregnenolone (**2**), consistent with its function as the sterol side chain cleaving enzyme (Fig. **3e,f**).

### *Using* cyp87a126 *mutant lines to investigate intermediates in cardiac glycoside biosynthesis*

*cyp87a126* mutant lines provide a tool for investigation of intermediates in cardiac glycoside biosynthesis, as only the first enzyme in the pathway is absent and the rest of the pathway remains intact. The following predicted intermediates were fed to *cyp87a126-2* plants: pregnenolone (**2**), isoprogesterone (**3**), progesterone (**4**), and 5β-pregnane-3,20-dione (**5**). All infiltrated substrates rescued cardiac glycoside biosynthesis in *cyp87a126-2* plants (Fig. **4**), consistent with the pathway shown in Fig. **1**.

**Fig. 4.**
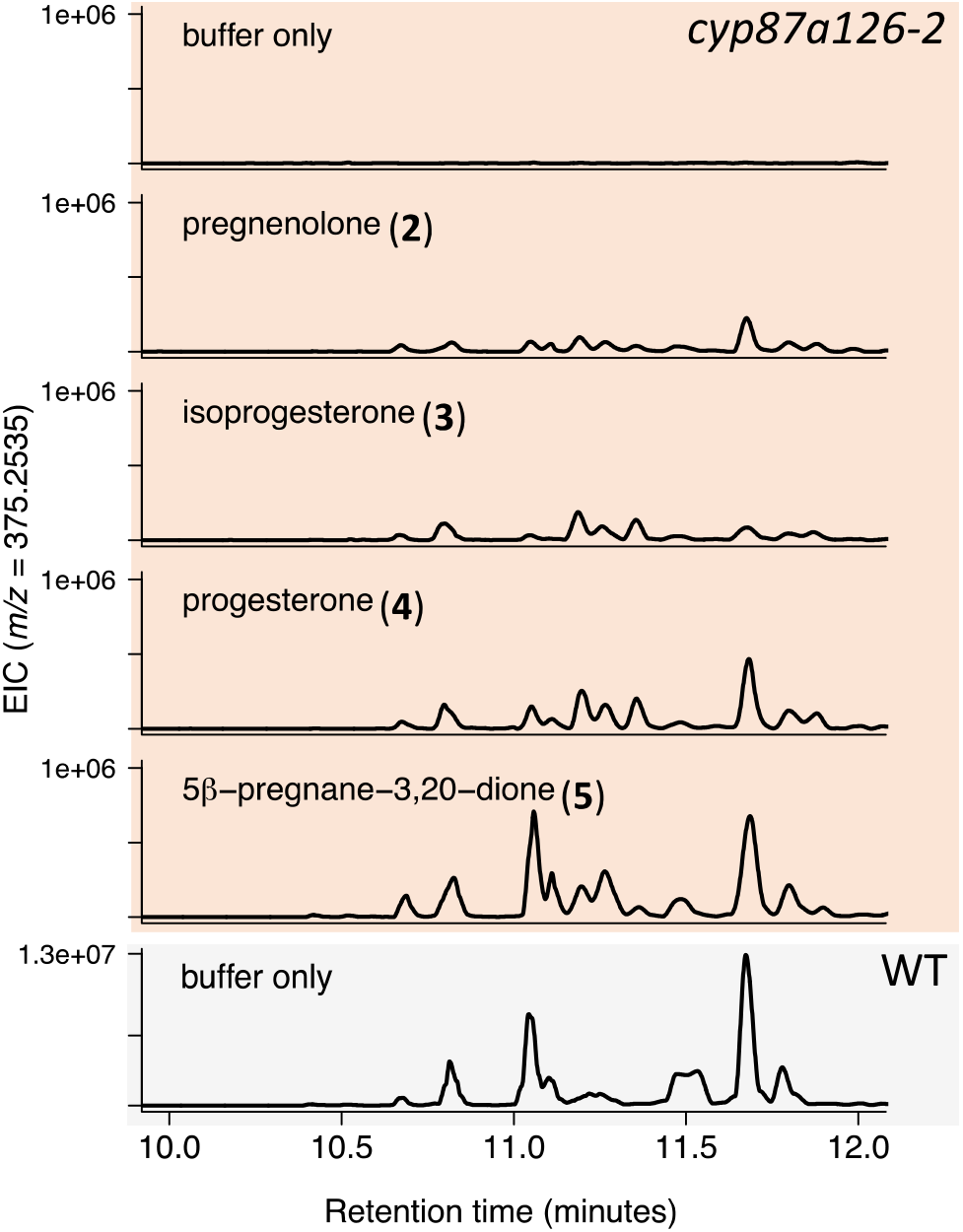
Rescue of cardiac glycoside biosynthesis in the *cyp87a126-2* mutant line. ESI+ extracted ion chromatograms of *m/z* = 375.2535, a fragment associated with digitoxigenin glycosides, from *cyp87a126-2* plants two days after substrate infiltration (orange background) compared to wildtype (grey background).

### *Leaf extracts of* cyp87a126 *mutant lines display decreased Na^+^,K^+^-ATPase inhibition*

Cardiac glycosides are strong inhibitors of Na^+^,K^+^-ATPase activity in animals (Petschenka *et al*., 2023). Therefore, we assessed the *cyp87a126* mutant lines for Na^+^,K^+^-ATPase inhibitory activity using an *in vitro* assay with porcine Na^+^,K^+^-ATPase. Methanolic extracts of *cyp87a126-1* displayed on average 15-fold lower inhibitory activity than extracts of wildtype leaves, and *cyp87a126-2* showed 251-fold lower inhibitory activity, on par with the cardiac glycoside-free *A. thaliana* control (Fig. **5a,b**, Table **S8**). A one-way ANOVA on log-transformed half maximal inhibitory concentration (IC50) data found differences between wildtype and both *cyp87a126* lines (one-way ANOVA, F_3,12_=69.34, *P<*0.001; Tukey’s HSD: WT-*cyp87a126-1 P*<0.001, WT-*cyp87a126-2 P* <0.001, *cyp87a126-1-cyp87a126-2 P* =0.051). Although we see some apparent Na^+^,K^+^-inhibition at high concentrations of both *A. thaliana* and cardiac glycoside-free *E. cheiranthoides* lines, this is an artefact of the assay which shows inhibitions at these concentrations with other cardenolide free plant extracts.

**Fig. 5.**
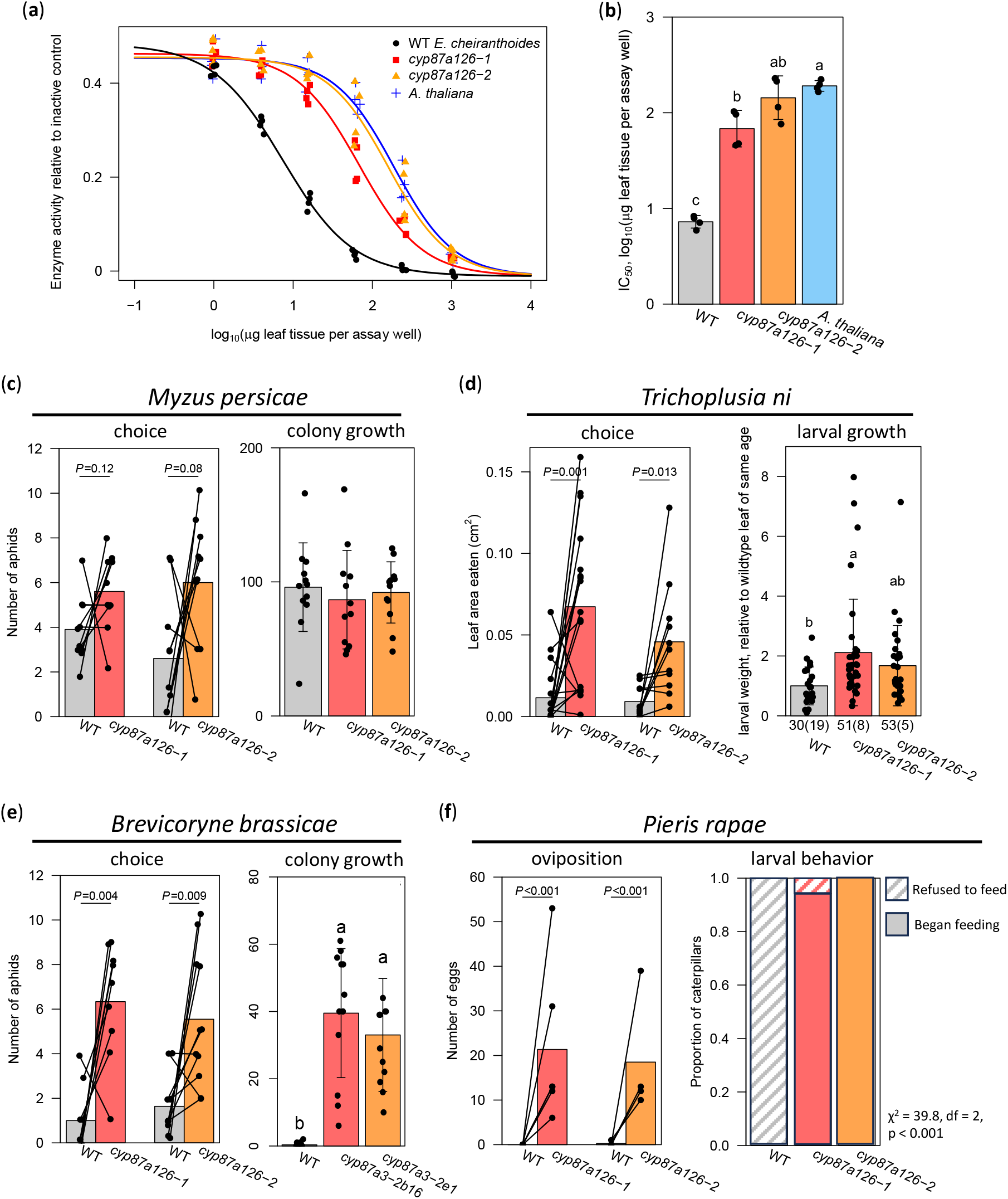
Functional implications of *cyp87a126* knockout and loss of cardiac glycosides. (a) Na^+^,K^+^ ATPase inhibition assay for leaf extracts of *cyp87a126* mutant lines compared with wildtype (WT) *E. cheiranthoides* and *A. thaliana* Col-0 as a cardiac glycoside-free control. Inhibition curves were calculated from four replicates of each tissue. (b) Half-maximal inhibitory concentration of leaf extracts estimated from inflection point of inhibition curves (n=4 plants per line). (c,e) *M. persicae* and *B. brassicae* aphid assays: binary choice as measured by aphid position after 24 hours (n=10 aphids per replicate, 9-11 replicates per line); colony growth of five synchronized aphids after 9 days (n=12 plants per line). (d) *T. ni* assays: binary choice, leaf area eaten after two days (n=18); growth, larval weight after 12 days of feeding, normalized by leaf position to remove effect of leaf age (n=25-37 caterpillars per line). Numbers below plot indicate caterpillars surviving or dying (in parentheses) after 8 days. (f) *P. rapae* assays: oviposition in binary choice assay (n=75-128 eggs); larval behavior when confined to an individual leaf (n=13- 17 caterpillars). For all plots: error bars indicate ± SD; letters indicate statistical differences (*P<*0.05) according to a one-way ANOVA with a post-hoc Tukey’s HSD, except for (e), where a Kruskal-Wallis test with pairwise Wilcoxon rank-sum comparisons. *P*-values are from paired t-tests in choice assays or chi-squared test in oviposition and larval behavior assays.

### *Insect performance on* cyp87a126 *mutant lines*

To test the impact of cardiac glycosides on insects feeding on *E. cheiranthoides*, we conducted insect choice and performance assays using two generalist herbivores, *M. persicae* and *T. ni*, and two crucifer-feeding specialists, *B. brassicae* and *P. rapae*. In choice assays, the overall trend was a preference for cardiac glycoside-free mutant lines, with varying levels of significance for each species. While more adult *M. persicae* aphids chose mutant lines over wildtype (*cyp87a126-1*: 56% chose mutant, *cyp87a126-2*: 67% chose mutant), this difference was not significant (Fig. **5c**, Table **S9**, paired *t*-test: *cyp87a126-1 P=*0.12, *cyp87a126-2*: *P=*0.08).

*Trichoplusia ni* caterpillars showed a clear preference for *cyp87a126* mutant lines over wildtype, as measured by leaf area eaten (Fig. **5d**, Fig. **S4**, Table **S10**, paired *t*-test: *cyp87a126-1 P=*0.001, *cyp87a126-2 P=*0.013). Similarly, adult *B. brassicae* aphids displayed a strong preference for cardiac glycoside-free mutant lines (Fig. **5e**, Table **S11**, paired *t*-test: *cyp87a126-1 P=*0.004, *cyp87a126-2*: *P=*0.009). Gravid adult *P. rapae* almost exclusively chose to oviposit on *cyp87a126* mutant plants, with all 128 eggs were laid on mutant plants for *cyp87a126-1* (χ^2^=128, df=1, *P<*0.001), and 74 of 75 eggs were laid on mutant plants for *cyp87a126-2*, with only one egg laid on wildtype (χ^2^=71.1, df=1, *P<*0.001) (Fig. **5f**, Table **S12**).

Results were less uniform in performance assays. For *M. persicae*, population growth over nine days from five adult aphids restricted to a single plant was not different between wildtype and either mutant line (Fig. **5c**, Table **S13**, one-way ANOVA: F_2,33_=0.27 *P=*0.77). When bagged on individual leaves, *T. ni* was more likely to refuse to feed on wildtype than on either mutant line (Fig. **5d**, χ^2^=17.44, df=2, *P<*0.001). Among caterpillars that did begin feeding, caterpillars grew marginally better on *cyp87a126-1* than on wildtype after correcting for leaf age, but there was no difference between *T. ni* growth on *cyp87a126-2* and wildtype (Fig. **5d**, Table **S14**, one-way ANOVA F_2,82_=4.18 *P=*0.019, Tukey’s HSD: WT-*cyp87a126-1 P=*0.014, WT-*cyp87a126-2 P=*0.22). Adult *B. brassicae* aphids were unable to establish colonies on wildtype plants, but five adult aphids grew to colonies averaging more than 30 aphids after nine days on both mutant lines (Fig. **5e**, Table **S15**, Krusal-Wallis, H_2_=24.37, *P<*0.001, Wilcoxon rank sum with Bonferroni correction: WT-*cyp87a126-1 P<*0.001, WT- *cyp87a126-2 P<*0.001, *cyp87a126-1*- *cyp87a126-2: P=*0.76). None of the 14 *P. rapae* caterpillars placed on wildtype plants began feeding, while 29 of 30 caterpillars placed on the two mutant lines fed and produced substantial damage (Fig. **5f**, Fig. **S5**, Table **S16**, χ^2^=39.8, df=2, *P<*0.001). While mortality of *P. rapae* caterpillars was high, four of those feeding on mutant plants reached pupation, demonstrating their suitability as a host plant.

### Herbivore attack in the field

On plants growing in the field, a total of 2,435 visitors from 30 distinct functional or taxonomic groups were recorded on *E. cheiranthoides* wildtype, *cyp87a126-1*, and *cyp87a126-2* across four censuses spanning one month. Rare visitors, defined as appearing on fewer than 25 unique plants (24% of all plants) across the four censuses, were not subjected to statistical analysis. Common visitors included striped flea beetles (*Phyllotreta striolata* F.), turnip aphids (*Lipaphis erysimi* Kaltenbach), leafhoppers, snails, spiders, leaf miners, and *P. rapae* eggs. Snails of the family Succineidae were less likely to be found on wildtype plants (WT observation rate: 0.44, 95% CI 0.31-.058, *P=*0.41) than on *cyp87a126-1* plants (*cyp87a126-1* observation rate: 0.74, 95% CI 0.55-0.87, *P=*0.013) or *cyp87a126-2* plants (*cyp87a126-2* observation rate: 0.92, 95% CI 0.72- 0.98, *P<*0.001) (Fig. **6c**). Similarly, *P. rapae* eggs were more likely to be found on either mutant line than on wildtype plants (Fig. **6d**; WT observation rate: 0.04, 95% CI 0.01-0.14, *P<*0.001; *cyp87a126-1* observation rate: 0.48, 95% CI 0.30-0.66, *P<*0.001; *cyp87a126-2* observation rate: 0.45, 0.27-0.65, *P<*0.001). Visitation rates did not differ between wildtype and *cyp87a126* mutant lines for *P. striolata* (Fig. **6a**, one-way ANOVA: F_2,97_=2.23 *P=*0.11), *L. erysimi* (Fig. **6b**, WT observation rate: 0.32, 95% CI 0.21-0.46, *P=*0.01; *cyp87a126-1* observation rate: 0.48, 95% CI 0.30-0.66, *P=*0.18; *cyp87a126-2* observation rate: 0.21, 0.09-0.41, *P<*0.29), leafhoppers, leaf miners, or spiders. Complete visitation records are and results of logistic regressions for all common taxa are in Table **S17** and Table **S18**, respectively.

**Fig. 6.**
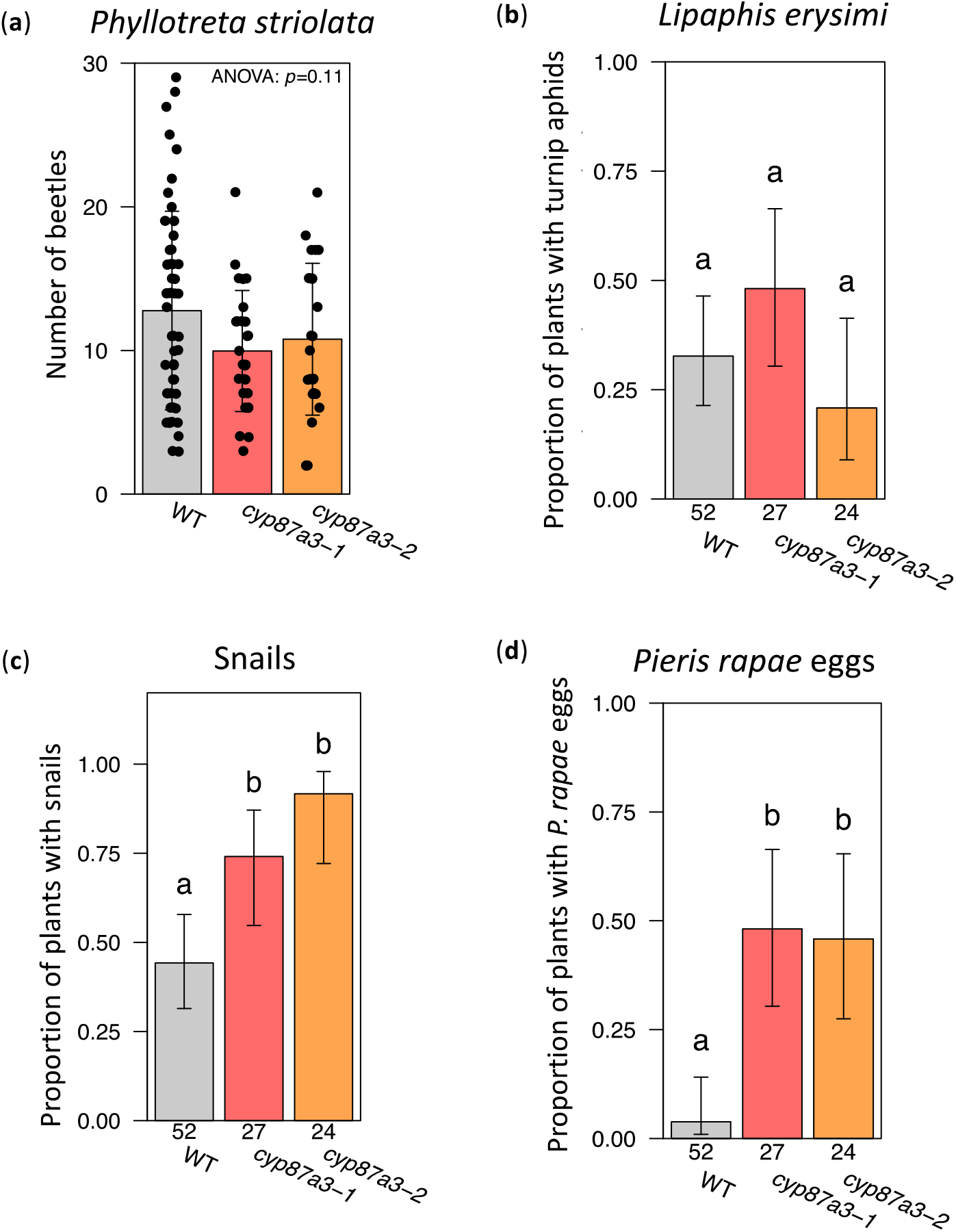
Observation rates of common visitors across four censuses in a field setting. (a) Count of flea beetles (*Phyllotreta striolata*) on each plant during the fourth census, when they were most abundant. Error bars indicate ± SD. Proportion of plants with (b) turnip aphid *(Lipaphis erysimi*), (c) snails of the family Succineidae, or (d) cabbage butterfly (*Pieris rapae*) eggs at any point across the four censuses. Numbers below bars indicate the number of surviving plants belonging to each genotype that were included in the analysis. For (b), (c), and (d), error bars indicate 95% confidence interval from logistic regression, and letters denote differences between estimated marginal means for each group, Tukey-adjusted *P*<0.05.

## DISCUSSION

### Convergent evolution of CYP87A126 as a sterol side chain cleaving enzyme and the first committed step in cardiac glycoside biosynthesis

The presence of sterol side-chain cleaving enzymes in cardiac glycoside-producing plants has been the subject of speculation for decades (Stohs & El-Olemy, 1971; Pilgrim, 1972; Lindemann & Luckner, 1997; Iino *et al*., 2007; Lindemann, 2015), and they have only recently been identified in *Digitalis* spp., *Calotropis procera*, and *E. cheiranthoides* (Carroll *et al*., 2023; Kunert *et al*., 2023). In this study, we independently confirm that *Ec*CYP87A126 possesses sterol side chain cleaving activity and generate knockout lines showing that this activity is required for cardiac glycoside production *in planta*. This discovery is an important first step towards establishing the full cardiac glycoside biosynthetic pathway in *Erysimum*. Based on and substrate feeding experiments, it is now clear that *Erysimum* cardiac glycoside biosynthesis proceeds through pregnane intermediates, much like in *Digitalis* (Kunert *et al*., 2023). Notably, the two amino acid substitutions identified by Carroll et al. (Carroll *et al*., 2023) that are necessary for sterol side chain cleaving activity in *Dl*CYP87A4, V355A and A357L, are also present in *Ec*CYP87A126. While the ancestral function of the CYP87A clade is unknown, other related enzymes from the CYP87 family are known to act on triterpenoids (Zhou *et al*., 2016; Ghosh, 2017).

### Identification of other cytochrome P450 monooxygenases involved in cardiac glycoside biosynthesis

Our screen of cytochrome P450 monooxygenases revealed a second P450 that is involved in cardiac glycoside modification. *Ec*CYP716A418 mutant lines still make high quantities of cardiac glycosides, but they almost exclusively accumulate digitoxigenin glycosides, which are not oxygenated at carbons 4 and 19 (Fig. **1**). Based on the predicted pathway, we hypothesize that *Ec*CYP716A418 hydroxylates digitoxigenin at carbon 19. However, we did not see this activity when co-infiltrating *Ec*CYP716A418 with digitoxigenin or digitoxin in *N. benthamiana*. This result leaves open the possibility that hydroxylation by *Ec*CYP716A418 occurs earlier in the pathway, for example prior to lactone ring formation. It is also possible that the observed phenotype is somewhat more cryptic, and there is not a direct link between this enzyme and cardiac glycoside hydroxylation. Other members of the CYP716A family are well known for the modification of triterpenoid scaffolds, including β-amyrin (Carelli *et al*., 2011; Ghosh, 2017). In addition, *Ec*CYP716A418 is duplicated several times in *E. cheiranthoides* relative to *A. thaliana* (Fig. **2d**), a pattern that is often observed in the evolution of specialized metabolic pathways (Moghe & Last, 2015).

A knockout line of the third cytochrome P450 discussed in this paper, *Ec*CYP71B132, did not have an altered cardiac glycoside phenotype. The lack of a phenotype does not conclusively exclude involvement of *Ec*CYP71B132 in cardiac glycoside biosynthesis. For example, a potential alternative start codon 50 base pairs after the Cas9-induced deletion may result in a functional protein with an *N*-terminal truncation of 66 amino acids, leaving the active site intact. Even if *cyp71b132-1* is a complete functional knockout, it is possible that its role in cardiac glycoside biosynthesis is complemented by a functionally redundant enzyme.

Nonetheless, the lack of known CYP71B family members acting on steroid-like compounds indicates that the involvement of *Ec*CYP71B132 in cardiac glycoside biosynthesis is less likely. The most closely related characterized enzyme, *At*CYP71B15, acts on indolic intermediates in camalexin biosynthesis (Böttcher *et al*., 2009).

### Cardiac glycosides as escape from herbivory

While it has long been understood that *Erysimum* represents a unique instance of the co-occurrence of two potent defensive compounds (Züst *et al*., 2018), the overall benefit to the plants from investing in two distinct but potentially redundant defenses has been difficult to test. In combination with the field experiment, our insect feeding and performance assays highlight that plant-insect relationships can be highly species-specific. In the laboratory, generalist *M. persicae* aphids performed similarly on *E. cheiranthoides* regardless of the presence of cardiac glycosides, and generalist *T. ni* caterpillars preferred the cardiac glycoside-free mutant but grew only slightly better when feeding on it. This is perhaps unsurprising as generalist insects are known for their ability to tolerate a wide range of defensive metabolites. Conversely, we found that snails from the family Succineidae, which are also broad generalists, were more likely to be found on cardiac glycoside-free mutant plants in the field. Although we did not quantify damage caused by snails, this is strong evidence that cardiac glycosides serve as a feeding deterrent against snails, consistent with previous work showing that cardiac glycosides can be effective molluscicides (Dai *et al*., 2011).

A much clearer impact of the loss of cardiac glycosides is the reversal of *Erysimum*’s escape from herbivory by two glucosinolate-feeding specialist herbivores. Consistent with previous reports (Rothschild *et al*., 1988; Renwick *et al*., 1989; Sachdev-Gupta *et al*., 1993), we found that both *P. rapae* and *B. brassicae* are unable to utilize wildtype *E. cheiranthoides* as a host plant. However, with the loss of cardiac glycoside biosynthesis in the *cyp87a126* mutant lines, *P. rapae* found *E. cheiranthoides* to be an acceptable host in both field and lab settings, and *B. brassicae* aphids established robust colonies in the lab. This greater sensitivity of these crucifer specialists to cardiac glycosides is consistent with a theory proposed by Cornell and Hawkins (Cornell & Hawkins, 2003), who concluded that specialist insect herbivores are less likely to tolerate novel defensive compounds like cardiac glycosides in *Erysimum*. However, two crucifer specialists observed in the field, *P. striolata* and *L. erysimi*, did not show a preference for cardiac glycoside-free *E. cheiranthoides*.

Complete escape from even a subset of specialist insects would represent a distinct ecological advantage, as specialist insects are observed to cause the majority of damage a plant suffers in certain contexts (Coley & Barone, 1996; Bidart-Bouzat & Kliebenstein, 2008). Despite this apparently clear defensive advantage, cardiac glycoside production has been lost or drastically reduced in the accession of *E. collinum* screened in this study (Züst *et al*., 2020). Whether this loss has become fixed would require more thorough sampling in its native range in Iran, but the persistence of even some individuals with a complete lack of cardiac glycosides points to context-dependent benefits and likely substantial costs of cardiac glycoside production. *Erysimum collinum* also accumulates high levels of glucoerypestrin, a glucosinolate unique to *Erysimum* (Kjær & Gmelin, 1957; Fahey *et al*., 2001; Blažević *et al*., 2020; Züst *et al*., 2020) that may have allowed an alternative escape route from glucosinolate specialists, rendering cardiac glycosides unnecessary as a second line of defense. By contrast, there are no known cases of the loss of glucosinolates in *Erysimum*, perhaps because glucosinolates are involved in non-defensive processes such as signaling and development (Katz *et al*., 2015). Future experiments including more extensive field experiments with both glucosinolate and cardiac glycoside knockouts will more fully dissect the complex role cardiac glycosides play in allowing *Erysimum* to escape herbivory and determining plant fitness in varied contexts.

## ACKNOWLEDGEMENTS

We thank Prof. Tobias Züst for his insightful comments on the manuscript, Prof. David Nelson for cytochrome P450 naming, Julia Dahl for her assistance with insect experiments, and Dr. Boaz Negin for assistance with the field trial. This research was funded by United States Department of Agriculture award 2020-60713-33565 and an award from the Triad Foundation to GJ; a Chemistry Biology Interface Training Program fellowship under National Institutes of Health/National Institute of General Medical Sciences (T32GM138826) and a US National Science Foundation Graduate Research Fellowship (DGE–2139899) to GCY; a Summer Undergraduate Research Fellowship from the American Society of Plant Biologists and a Rawlings Cornell Presidential Research Scholar award to MLA; and a fellowship from the Cane-Bridge Foundation to APC.

## AUTHOR CONTRIBUTIONS

GCY, MLA, HDF, MM, and GJ designed the research; GCY, MLA, HDF, MM, APC, and APH performed the research; GCY and MLA analyzed data; GCY, AAA, and GJ wrote and edited the manuscript.

## DATA AVAILABILITY

The data that support the findings of this study are available in the Supporting Information. Raw sequencing reads are publicly available on NCBI (PRJNA1015726). Seeds from mutant lines are available from the Arabidopsis Biological Resource Center (https://abrc.osu.edu/).

## SUPPORTING INFORMATION

**Fig. S1** Maps for plasmids used to generate Cas9 constructs.

**Fig. S2** Experimental setup for insect assays.

**Fig. S3** Gene models for Cas9 targets and knockout lines.

**Fig. S4** Photos of leaf disks from *Trichoplusia ni* choice assay.

**Fig. S5** Photos of leaves from *Pieris rapae* feeding assay.

**Table S1** Sequences for primers used in this study.

**Table S2** *m/z* values and retention times used for quantifying cardiac glycosides in LCMS data.

**Table S3** *m/z* values and retention times used for quantifying glucosinolates in LCMS data.

**Table S4** Full length coding sequences for cytochrome P450s.

**Table S5** Relative peak area for *cyp716a418* cardiac glycosides.

**Table S6** Relative peak area for *cyp87a126* cardiac glycosides.

**Table S7** Relative peak area for *cyp87a126* glucosinolates.

**Table S8** Raw data from Na^+^,K^+^-ATPase assay.

**Table S9** Raw data from *Myzus persicae* choice assay.

**Table S10** Raw data from *Trichoplusia ni* choice assay.

**Table S11** Raw data from *Brevicoryne brassicae* choice assay.

**Table S12** Raw data from *Pieris rapae* oviposition assay.

**Table S13** Raw data from *Myzus persicae* growth assay.

**Table S14** Raw data from *Trichoplusia ni* growth assay.

**Table S15** Raw data from *Brevicoryne brassicae* growth assay.

**Table S16** Raw data from *Pieris rapae* feeding assay.

**Table S17** Complete observation records from field experiment.

**Table S18** Logistic regressions for common taxa in field experiment.

