## Supplemental Figures and Tables for "Cardiac glycosides protect wormseed wallflower (*Erysimum cheiranthoides*) against some, but not all, glucosinolate-adapted herbivores"

***New Phytologist* Supporting Information**

The following Supporting Information is available for this article:

**Fig. S1** Maps for plasmids used to generate Cas9 constructs.

**Fig. S2** Experimental setup for insect assays.

**Fig. S3** Gene models for Cas9 targets and knockout lines.

**Fig. S4** Photos of leaf disks from *T. ni* choice assay.

**Fig. S5** Photos of leaves from *P. rapae* feeding assay.

**Table S1** Sequences for primers used in this study.

**Table S2** *m/z* values and retention times used for quantifying cardiac glycosides in LCMS data.

**Table S3** *m/z* values and retention times used for quantifying glucosinolates in LCMS data.

**Table S4** Full length coding sequences for cytochrome P450s.

**Table S5** Relative peak area for *cyp716a418* cardiac glycosides.

**Table S6** Relative peak area for *cyp87a126* cardiac glycosides.

**Table S7** Relative peak area for *cyp87a126* glucosinolates.

**Table S8** Raw data from $Na^+$,$K^+$-ATPase assay.

**Table S9** Raw data from *Myzus persicae* choice assay.

**Table S10** Raw data from *Trichoplusia ni* choice assay.

**Table S11** Raw data from *Brevicoryne brassicae* choice assay.

**Table S12** Raw data from *Pieris rapae* oviposition assay.

**Table S13** Raw data from *Myzus persicae* growth assay.

**Table S14** Raw data from *Trichoplusia ni* growth assay.

**Table S15** Raw data from *Brevicoryne brassicae* growth assay.

**Table S16** Raw data from *Pieris rapae* feeding assay.

**Table S17** Complete observation records from field experiment.

**Table S18** Logistic regressions for common taxa in field experiment.

**Fig. S1.** Plasmid maps for pARV370, pARV380, and pARV483 used for generating Cas9/gRNA constructs.

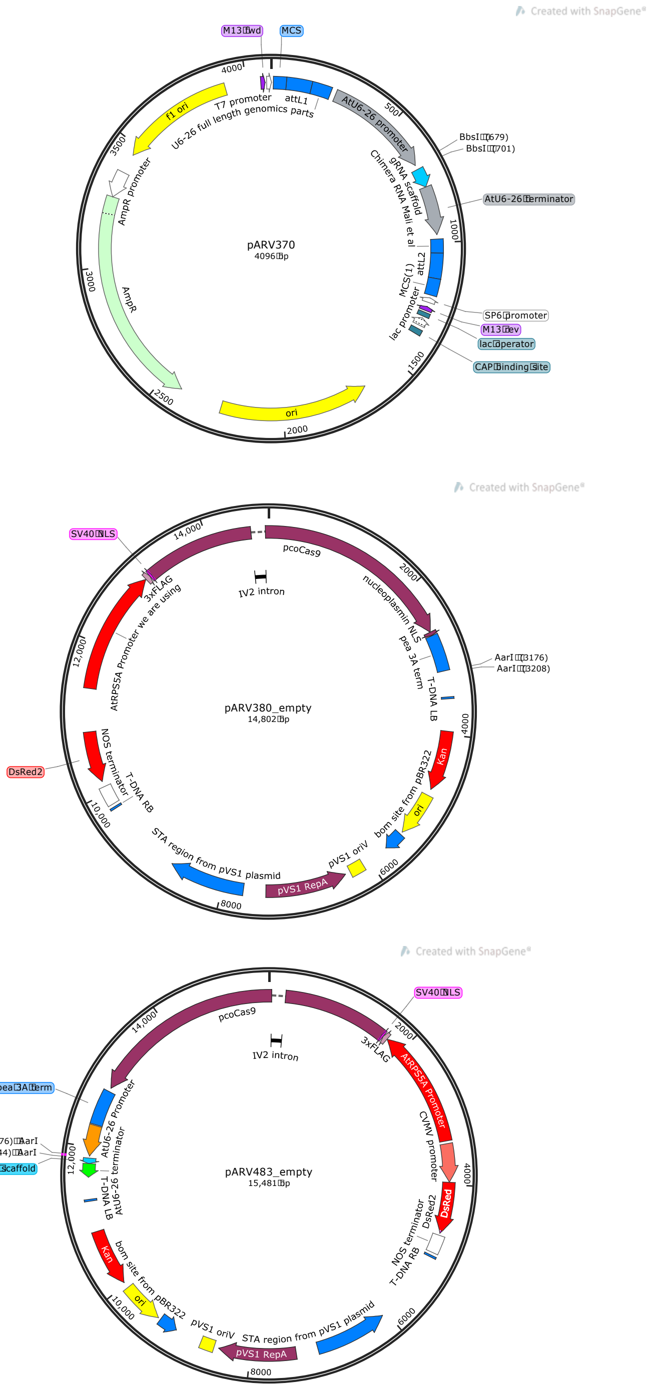

**Fig. S2**. Experimental setup for insect assays. (A) *T. ni* larval choice. Neonate larvae were placed between the two leaf discs. (B) *M. persicae* choice. Ten synchronized adult aphids were placed in each Petri dish between the leaf tips. (C) *P. rapae* feeding. A single neonate larva was placed in each bag, one bag per plant. The setup for the *T. ni* feeding experiment was similar, but there were five bags per plant. (D) *P. rapae* oviposition. One male and one female *P. rapae* butterfly were placed in a cage with one wildtype and one *cyp87a126* mutant plant.

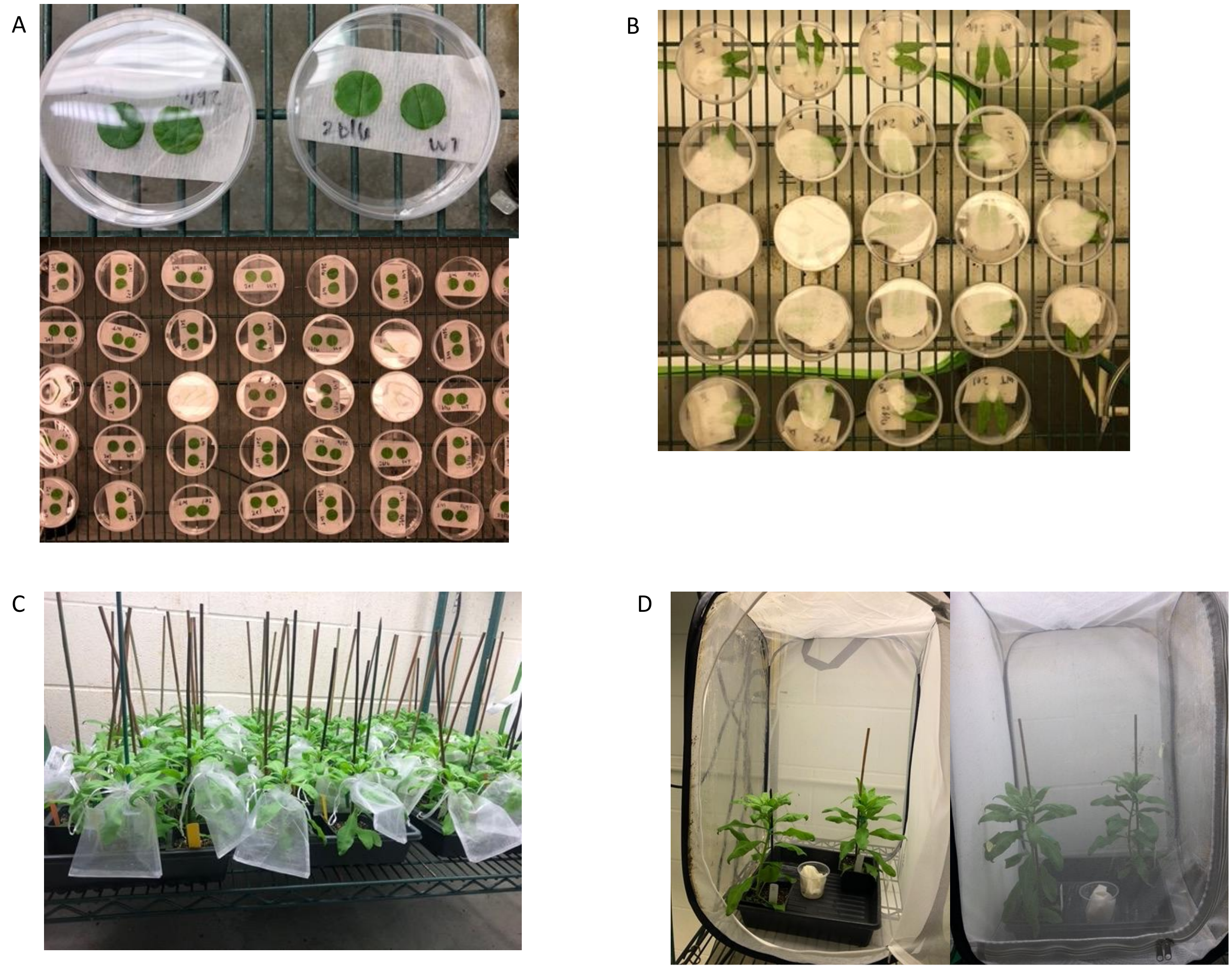

**Fig. S3**. Graphical representation of all three cytochrome P450s investigated in this paper in both wildtype and mutant lines. Untranslated regions are shown in light blue, coding regions in dark blue, and introns as thin lines. Regions translated out of frame and premature stop codons in mutant lines are shown in red.

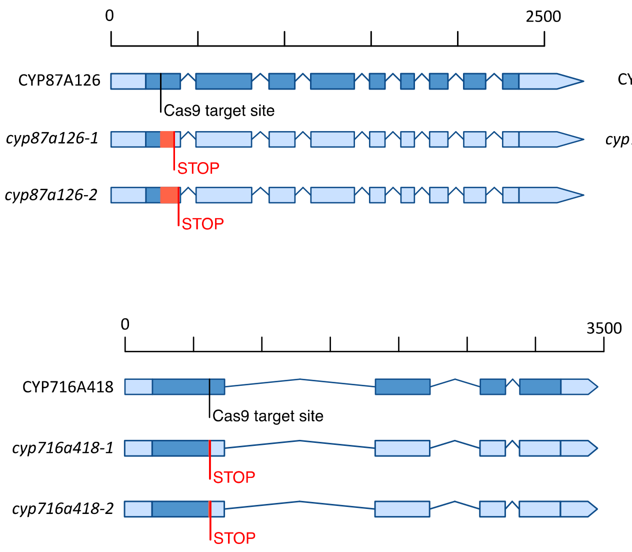

**Fig. S4**. Leaf discs from *T. ni* choice assay after 48 hours of feeding. Paired discs from the same Petri dish are next to each other. Total leaf area consumed was quantified from these photos using LeafByte. Dots are 4.8 cm apart.

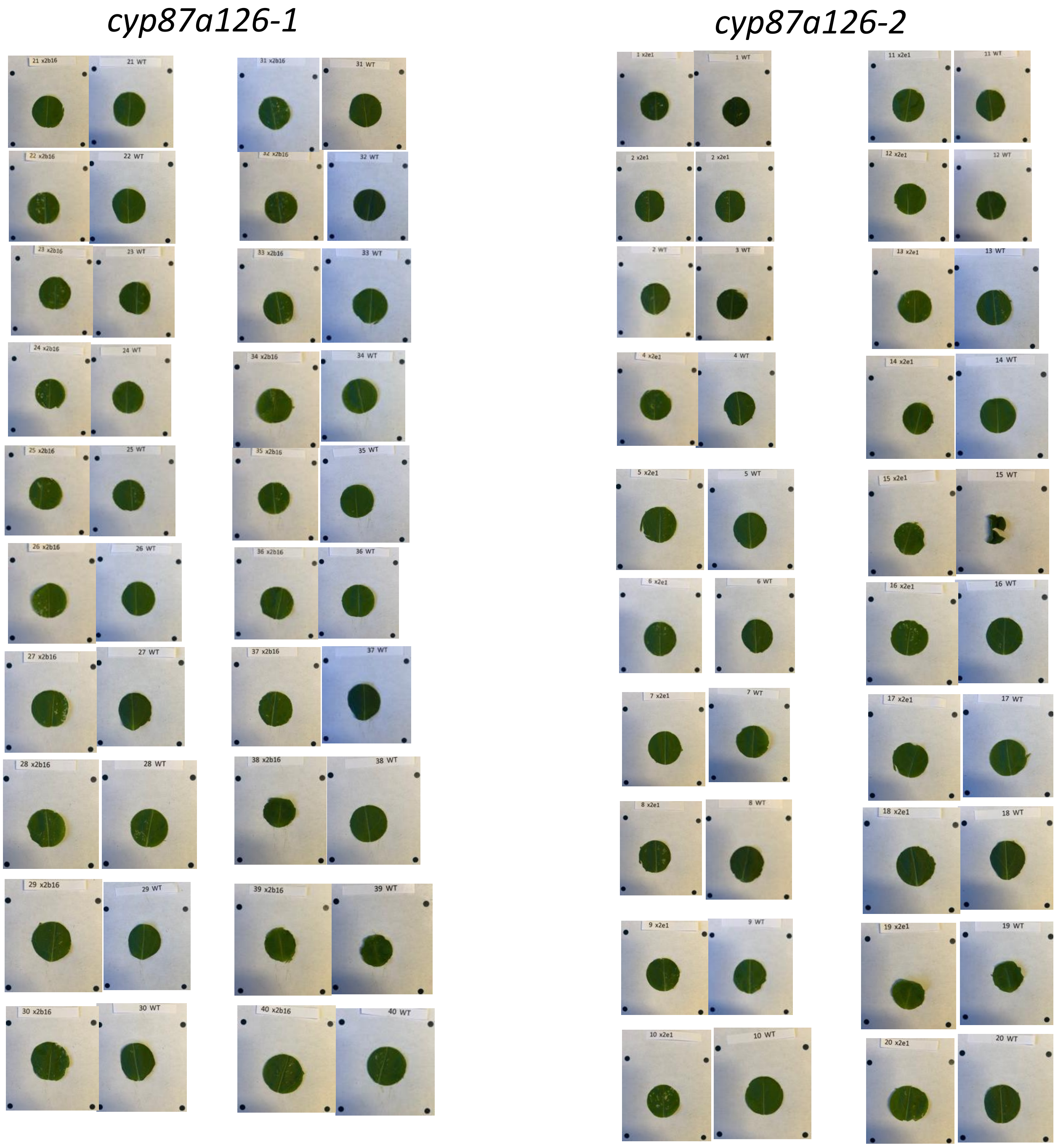

**Fig. S5**. Leaves from *P. rapae* feeding assay after 8 days of feeding. At this point, caterpillars had either died or been moved to a fresh leaf. Leaves were scored as either “damaged”, or “undamaged.” Distance between dots is 4.8 cm.

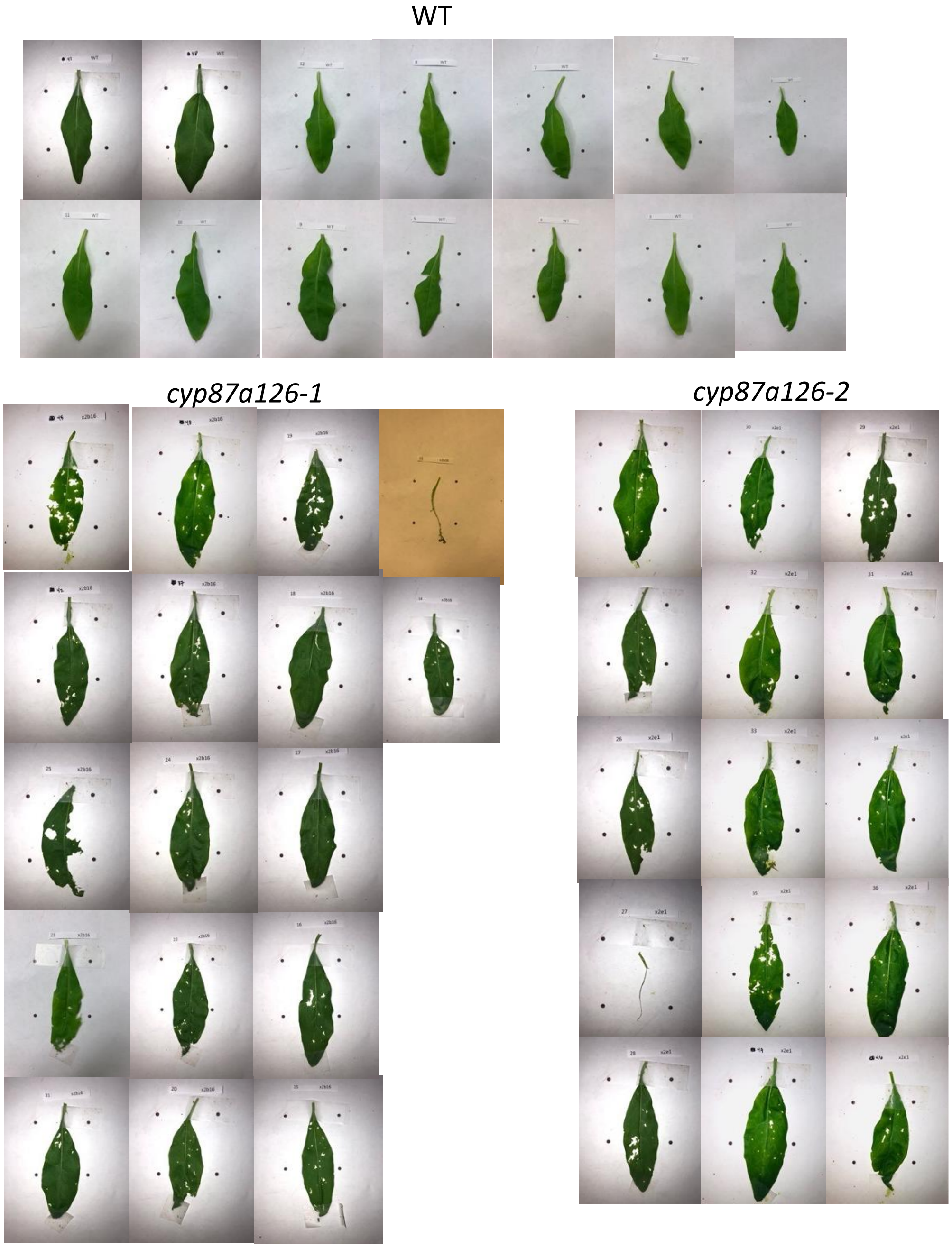

**Table S1** Sequences for primers used in this study.

See Excel spreadsheet.

**Table S2** *m/z* values and retention times used for quantifying cardiac glycosides in LCMS data.

See Excel spreadsheet.

**Table S3** *m/z* values and retention times used for quantifying glucosinolates in LCMS data.

See Excel spreadsheet.

**Table S4**. Full length coding sequences for the three cytochrome P450s discussed in this study.

```
>Erysimum_cheiranthoides_CYP87A126
ATGTCGTGGGCATTATGCATTTGGGTTTCTCTAGTTGTGACAGGCATCACAACTTTGGTT
TACAAATGGAGGAACCCTAAATGCAGTGGGAAGCTTCCACCAGGTTCTATGGGCTTACCG
CTTCTGGGCGAGACGATTCAATTCTTCAAACCAAACTTAACTTCAGACATCCAACCCTTT
ATTAAAGAGAGGACTAAGAAGTATGGGCCAATCTTCAAGACCAGCCTTGTGGGGAAATCA
ATTATTGTAACCACAGACCCTGATTTTAGTTATTTTGTGTTCCAACAAGAAGGTCAATCT
TTCCAGAGTTGGTATCCTGACACTTTCGTTGAGATCTTTGGAAAGCAGAATTTGGGAGCA
TTGCATGGGATCATTTATAAGTACCTCAAGCACATGGTCTTGAGTCTCTTTGGCTTTGAA
AGTCTCAAGAATATGCTTCCTGAAATTGAACAAACTGCATGTAAGAAGCTAGACCTCTGG
TCAACTCAAAAATCAATTGAGCTAAAAGAGTCCACGGCAAACTTGATATTTGATCTCACC
GCGAAAAAGTTGATTAGCCATGATGAAGAGAAGTCCTCAGAGAATCTAAGGGACAACTAT
GTTGCTTTCATCGATGGTTTGATCTCCTTCCCTATTAACCTCCCAGGGACAGCTTTTTAC
AAGTGCCTAAAGGGTAGGGAAAGGGTGATGTCAAGTTTGAGGAATATGCTTAAGGAGAGG
CGTAAGAATCCAAGGAAGGTTGCAAGTGACTTCTTCGATTATGTCATCGAAGAGCTTAAG
AAAGAAGGGACAATGCTGACAGAATCAATTGCACTAGACTTGATGTTTGTCTTGCTTTTC
GCCAGCTTTGAAACAACATCTCTCGCTATTACTGTAGCTATCAAGATGCTCTCAGACCAC
CCTTCAGTCCTAAAACGTCTAACGGAAGAACATGAGGCAATTCTGAGAAACCGTAAAGAT
CCCAACTCTGGACTGACATGGGAAGAATACAAATCAATGACTTACACATTTCAGTTCATG
AACGAAACAGCGAGACTAGCGAATATAGCTCCTTTAATCTGCAGGAAAGCCTTGAAAGAT
ATCCAGTATAAAAATTATACCATTCCAGCCAATTGGCCAGTGATGGTTGTTCCTCCAGCT
ATACATTTGGATCCAAACAACTATGAGGATCCACTTGTCTTCAATCCATCTAGATGGGAA
GGATCAGAGTTTACTAATGCATCAAAGAAGTTCATGGCGTTTGGTGGCGGAATGCGATTT
TGTATCGGAACTGACTTTAGCAAACTCCAAACCGCAGTTTTCCTTCATAGCTTGCTCACA
AAATACAGTTGGGAGCATATCAGTGGAGGAAATATGCTTCGAAGTCCTGGATTGCAATTT
CCAAATGGGTATCATGTCAAAATCAACAAGAAAGAGATATGA
>Erysimum_cheiranthoides_CYP71B132
ATGGCTACCATTTGGATTCTTCTATCACTTCTCTTCCTAATCATCTTTCTTCTTCTCGCC
GCCTTCAGACGCACAAATCAGGGGAAACATCGACCAATACCATCTCCTCCTGGTTTTCCG
ATTATCGGAAACTTACACCAGCTTGGACAATTACAACATCAGTCTTTATGGAATCTCTCA
AAGAAGTATGGTCCTGTGATGCTTTTGAAGCTTGGAAAAGTACCAACAGTCATACTTTCT
TCCTCCGAAACAGCAAGACAAGCTCTACGAGAGAATGACCTCCATTGTTGTAGCCGTCCT
CTCTCAGCAGGGACAAGAGAGCTCTCTTACAACAATCTAGATATGTCTTCCTCTCCTTAT
AATGAGTACTGGAAAGAACTAAGGAAGCTCTGTGCTCAAGAACTCTTCAGTGCTAAAAAA
ATTCAATCCATTCAACCCATTAAGGACGAGGAGGTCAAGAAAGTAATCGATTCAATCGCT
GAGTCAGCGTCTCTGAAAAATCCGGTTAACTTGAGCACGAAGTTTCTTGCTTTAACAACA
TGCGTAGTTTGCAAGGCAGCATTAGGTGTGAGTTTTGAGGGAACTGTGCTCAACAGTGAC
AGATTCAACAAGTTAGTCCGAGATGCTTTCGAGATGTTGGGAAGCTTCCCTGCCTCAGAT
```

```
TTTATTCCGTATGTCGGATGGATCATCGATCACTTCCAAGGTTTACAAGGGATGAGAGAC
AAAAGTAAGCGAGATCTCGATGCGTTCTATGAACAAATGTTTGATCTGCATTATAAGGAA
AAAGAAGAAGGGAGTGAAGACTTCGTGGATCTTCTCTTGAGGTTGGAGAGAGAAGAAATT
GTTGTTGGAGATGCCAAGCTCACAAGAAATCATATCAAAGCAATTTTGATGAACATTCTT
TTAGGAGGAATCGATACTTCTGCAATCTCAATGACATGGGCAATGGCAGAACTTGCTAAG
AACCCAAGAGTGATGAAAAAAGTTCAATCCGAAATCAGAAGCCAAATCAAGAACAAAGAA
AGAATCAGCTTCGATGACACTGATAAGCTCGAGTACCTGAAAATGGTGATCAAAGAAACA
TGGAGGTTACATCCTCCAACACCTCTTCTGCTCCCAAGAGAAGTAACGACTGAATTCGAG
ATCAATGGCTACACAATTCCAGCCAAGACACGGCTTCATGTGAATGTATGGGCTATTGGG
CGTGATCCCGATACCTGGAAAGATCCAGAAATGTTTTTTCCTGAAAGGTTTATGGATAGT
AACATTGATGCAAAAGGACAGGACTTTGAGCTGTTATCGTTTGGGAGTGGTAGGAGAATA
TGTCCTGGAATGTACATGGGAGCAACAATGGTGGAGTGTGGCCTAGCTAACATGTTGTAT
CATTTTGATTGGAAATTACCAGAAGGCATGGCAGTCGAAGATATCGATATGGAAGAATCT
CCTGGACTTACTATAAGCAAAAAAAATGATCTTCTACTTGTTCCTGTGAAGTATTTGGAT
CACTGA
>ErysimuM_cheiranthoides_CYP716A418
ATGTATTTGACAATAATATTCCTCTTCGTTAGCTTCTTTCTTCTTCCTCTCTTGTTTCTCTTTGGAAAACATTT
ATCAAACATCTGTTATCGCAACCTTCCTCCGGGAAAGATTGGCTTTCCCTTAACCGGAGAGACTTTTTCCTTCC
TCTCTGCCGGCCGTCAAGGCCACCCGGAGAAGTTTATCACCGACCGAGTTCGTCGTTTCTCCTCAGGTGTCTTC
AAGACCCACCTCTTTGGGTCTCCCTTCGCGGTGGTGACCGGTGCGTCCGGGAACAAGTTTCTATTCACCAACGA
GAGCAAGCTCGTGGTCTCATGGTGGCCAGATTCCGTTACTAAGATCTTCCCTTTTCCAACGGAGTCAAACCCCA
CGGATGAGGCTCTTAGGACGAGGATACTTCTTGCGCCGGCCATGAAGTCTGAGGCTTTGAGGAACTATGTAGGT
ATTATGGATGAGGTGGCTCAGAGACAATTTGATACGGATTGGGCCAATCAAGATCAACTTATTGTCTTCCCTCT
CGCCCAAAAGTTCCAATTTGCTGTAGCATGCCGTTTGTTCATGAGTATGGACGACGAAGGAAGTGTAAGTAAAC
TCGAAAAGCCATTCAATGCAGTGTTAGCAGGGCTCGTCTCAATCCCAATAGATTTTCCAGGAACACGTTTTAAC
CGAGCCATTAAGGCATCGAGGTTAATTAGAAAAGAGGTTACCGCGATCATTAGGCAGAGAAAAGAAGAGCTGAA
GGCCGGGAAAGCATCATCGGAACAGGACATCCTATCTCACTTGTTGATAGGAGAGATCAAAGACGAGGATTTGG
CTATTAAGATTATTGCATTGTTAATGGCAGGACATGACACAACGAGTATCGTATTGACATTCATTGTCAACTAT
CTTGCTGAGTACCCTCATGTCTATCAACGTGTGTTACAAGAGCACAAGGATATACTAAAAGAAAAGGAAGGAAA
TGGAGCATTAACGTGGGAGGACATTTCGAAAATGAAATATTCTTGGAACGTTGCATGTGAAGTGATGCGAATTG
TTCCTCTTGCTGGCAGTTTTCGTGAGGCCATCAAAAGCTTTTCTTTCAAGGGTTTTAACATTCCCAAAGGATGG
AAGTTATATCGGAATGCCACCACAACACATATGAATCCAGAATACTTTCCGGAACCAGAGAAATTTGAGCCAAA
CCGATTTGAAGGAACTGTTCCGAAGCCTTATACATACGTTCCCTTTGGAGGAGGACCCAGGATCTGTCCAGGGA
GAGAGTATGCTCGTATAGAGATTCTTGTATTTATGCACCATCTTGTTAAGAGATTCGAGTGGGAAAAAGTGTTT
CCAGAAGAAAATAGACTAGTGGCTGACCCCAACCCAAAACCACCCAGAGGTCTCCCTATTAGAATTTTCCCTCA
ATCTGGATTTGAAATTTGA
```

**Table S5** Relative peak area for *cyp716a418* cardiac glycosides.

See Excel spreadsheet.

**Table S6** Relative peak area for *cyp87a126* cardiac glycosides.

See Excel spreadsheet.

**Table S7** Relative peak area for *cyp87a126* glucosinolates.

See Excel spreadsheet.

**Table S8** Raw data from $Na^+$,$K^+$-ATPase assay.

See Excel spreadsheet.

**Table S9** Raw data from *Myzus persicae* choice assay.

See Excel spreadsheet.

**Table S10** Raw data from *Trichoplusia ni* choice assay.

See Excel spreadsheet.

**Table S11** Raw data from *Brevicoryne brassicae* choice assay.

See Excel spreadsheet.

**Table S12** Raw data from *Pieris rapae* oviposition assay.

See Excel spreadsheet.

**Table S13** Raw data from *Myzus persicae* growth assay.

See Excel spreadsheet.

**Table S14** Raw data from *Trichoplusia ni* growth assay.

See Excel spreadsheet.

**Table S15** Raw data from *Brevicoryne brassicae* growth assay.

See Excel spreadsheet.

**Table S16** Raw data from *Pieris rapae* feeding assay.

See Excel spreadsheet.

**Table S17** Complete observation records from field experiment.

See Excel spreadsheet.

**Table S18** Logistic regressions for common taxa in field experiment.

See Excel spreadsheet.
